# In vivo gene disruption and homology-directed repair in muscles and muscle stem cells using CRISPR/Cas9

**DOI:** 10.64898/2026.06.30.735705

**Authors:** Bryan L. Peacker, Kuan-Hung Lin, Amy Lam, Christopher L. Rios, Kexian Zhu, Jill M. Goldstein, Kathleen Messemer, Regan Ellis, Michael Florea, Heidi Kletzien, Naftali Horwitz, August D. Bratti, Umika S. Paul, Madeline Maier, Medha KC, Tina Liu, Sara Ashrafi Kakhki, Ru Xiao, Luk Vandenberghe, Amy J. Wagers

## Abstract

Programmable endonucleases such as CRISPR/Cas9 provide powerful tools to edit mammalian genomes by engaging cellular mechanisms of DNA double-strand break (DSB) repair. CRISPR-catalysed homology-directed repair (CRISPR-HDR), though generally less efficient than other modes of DNA repair, holds particular promise to enable precise sequence replacement by targeted insertion of a homologous DNA template^1,2^. While recent studies have reported appreciable levels of HDR in cardiomyocytes in vivo^3^, skeletal muscle myofibres have historically been considered refractory to HDR-mediated genome editing^4^. Furthermore, how repair outcomes differ across tissues after systemic delivery of CRISPR/Cas9 editors, whether precise HDR editing can be achieved in regenerative tissue stem cells, and how developmental timing influences accessibility to CRISPR-induced repair remain unclear. Here, we use an adeno-associated virus (AAV)-delivered in vivo GFP-to-BFP colour-switching reporter system (AAV-GFP-to-BFP) to examine in vivo CRISPR-HDR with cellular- and tissue-level resolution. We find that postnatal cardiac muscle, skeletal muscle, and muscle stem cells undergo templated HDR at different rates across discrete developmental stages in mice. While HDR-edited muscle stem cells and myofibres were readily detectable after in vivo editing in juvenile mice, editing in neonatal mice yielded more efficient HDR in cardiac tissue. Based on these results, we adapted the CRISPR-HDR approach to rescue the therapeutically relevant *Dmd* mutation in *mdx* mice, demonstrating recoding to the wild-type protein sequence in both skeletal and cardiac muscles. These results provide a framework for advancing donor-templated DNA repair in living postnatal animals, and reveal unexpected cellular, developmental, and disease-related constraints on precise, therapeutic in vivo gene correction.

---

To sensitively detect in vivo CRISPR/Cas9 editing events, we designed a fluorescent reporter system (Fig. 1a), leveraging a transgenic mouse line that ubiquitously expresses enhanced green fluorescent protein (GFP)^5^. Using previously-described blue fluorescent protein (BFP) variants,^6–8^ we constructed a BFP HDR donor template carrying a minimal 2-base substitution (C197G and T199C) that creates a BtgI site and enables green-to-blue chromophore conversion. This modification simultaneously permits restriction fragment length polymorphism (RFLP) analysis and fluorescence-activated cell sorting (FACS) (Fig. S1). We also designed a *Staphylococcus aureus* Cas9 (SaCas9)-compatible guide RNA (gRNA) targeting the GFP chromophore region and found efficient disruption of GFP signal in GFP^+/-^;*mdx* tail tip fibroblasts (Fig. S1b,c). The promoterless BFP HDR donor was then combined with the GFP-gRNA in a single plasmid (GFP-gRNA-BFP-Template) for use in HDR experiments (Fig. 1b).

**Fig. 1.**
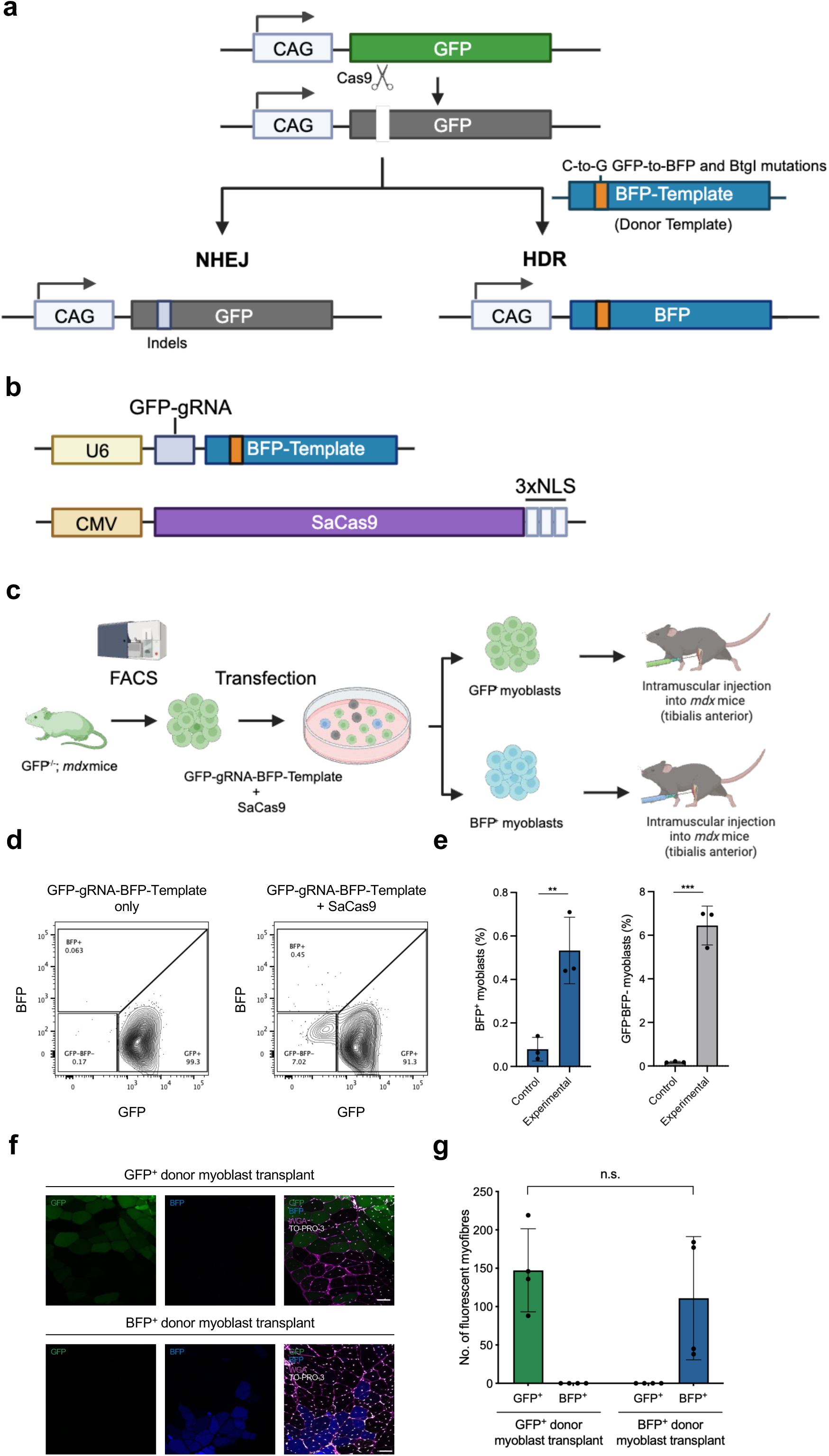
GFP/BFP colour switching reporter system enables discrimination and tracking of NHEJ- and HDR-edited myoblasts. **a**, Schematic of blue/green colour switching reporter for discriminating HDR vs. imprecise NHEJ. In the presence of a donor template, imprecise NHEJ disrupts GFP fluorescence while HDR-mediated substitutions enable GFP-to-BFP spectral conversion and create a BtgI restriction site for restriction fragment length polymorphism analysis. **b**, Plasmid constructs used for transfection and adeno-associated virus production. U6, U6 promoter; CMV, CMV promoter; NLS, nuclear localization signal. **c**, Experimental design. Skeletal muscle stem cells (satellite cells) were isolated from GFP^+/-^; *mdx* mice carrying a single transgenic CAG-GFP allele and transfected with plasmid constructs shown in **b**. Transfected cells were expanded in culture, then FACS-sorted on blue or green fluorescence for intramuscular transplantation into pre-injured recipient *mdx* mice. **d**, Representative flow cytometric analysis of myoblasts transfected with GFP-gRNA-BFP-Template alone or myoblasts transfected with SaCas9 and GFP-gRNA-BFP-Template. **e**, Frequency (%) of CRISPR-HDR edited BFP^+^ myoblasts and CRISPR-NHEJ edited GFP^-^BFP^-^ myoblasts in control or experimental cultures. Individual data points are shown overlaid with mean ± SD and represent n = 3 independent transfections. \*\**P* < .01,\*\*\**P* < .001, unpaired two-tailed t-test, **f**, Fluorescence detection of GFP^+^ and BFP^+^ myofibres in mice transplanted with sorted GFP^+^ (unedited) donor myoblasts or sorted BFP^+^ (HDR-edited) donor myoblasts. Scale bar, 50um. Green, GFP; Blue, BFP; Magenta, wheat germ agglutinin (WGA); White, TO-PRO-3. **g**, Quantification of fluorescent GFP and BFP myofibres in mice transplanted with GFP+ and BFP+ donor myoblasts. Data are presented as mean ± SD and represent n = 3 independent transfections. **e** and **g** were analysed using the Mann-Whitney *U* test. ** and *** indicate *P <* .01 *and P* < .005, respectively.

Using this colour-switching system, we first aimed to test the capacity of CRISPR/Cas9 to mediate HDR in vivo in a regenerative stem cell population – muscle satellite cells. Satellite cells isolated from skeletal muscles of GFP*^+/-^*;*mdx* mice were expanded ex vivo^9,10^ and transfected with plasmids containing SaCas9 and GFP-gRNA-BFP-Template, or GFP-gRNA-BFP-Template alone (Fig. 1b, c). Flow cytometry of cells transfected with both plasmids revealed a subset that lost green fluorescence (GFP^-^), indicative of imprecise NHEJ-mediated disruption of the GFP reading frame, as well as a separate population that both lost GFP and gained BFP signal (BFP^+^), indicative of HDR (Fig. 1d, e). In contrast, GFP^-^BFP^-^ and BFP^+^ cells were absent from cells receiving GFP-gRNA-BFP-Template alone (Fig. 1d, e). Blue fluorescence was absent when SaCas9 and gRNA were transfected alone, without the BFP-Template (Fig. S1d), indicating that NHEJ alone is unable to induce a GFP-to-BFP spectral shift. To assess NHEJ and HDR-mediated changes at the genomic level, we sorted the GFP^-^BFP^-^ and BFP^+^ populations separately by FACS and performed RFLP analysis and Sanger sequencing, which confirmed editing by CRISPR-NHEJ and -HDR, respectively (Fig. S2a, b). Finally, we investigated whether ex vivo edited satellite cell-derived myoblasts retained muscle-forming capacity. Upon inducing differentiation, both GFP^-^BFP^-^ and BFP^+^ satellite cell-derived myoblasts fused to form myosin heavy chain-positive myotubes (Fig. S2c). Furthermore, when transplanted into the pre-injured tibialis anterior muscle of *mdx* mice, sorted BFP^+^ myoblasts contributed to in vivo muscle repair, giving rise to BFP^+^ myofibres (Fig. 1f). Compared to muscle transplanted with GFP^+^ donor myoblasts, muscle transplanted with BFP^+^ donor myoblasts showed no difference in the number of fluorescent myofibres (Fig. 1g). These data indicate that the GFP-to-BFP colour-switching system reliably and sensitively reports genomic CRISPR-mediated NHEJ and HDR editing events with single-cell resolution and allows for subsequent assessment of the in vivo regenerative capacity of the edited cells.

We next evaluated whether the GFP-to-BFP reporter system could track CRISPR-mediated genome editing in vivo, delivering both CRISPR/Cas9 and template DNA via adeno-associated virus (AAV) vectors. Due to the limited cargo capacity of AAVs, we implemented a dual-vector strategy in which one vector encoded SaCas9 (AAV-SaCas9) and another encoded the GFP-gRNA-BFP-Template cassette (AAV-GFP-gT; Fig. 2a). We selected AAV8 because of its high liver, heart, and skeletal muscle tropism^11^. Juvenile (P21) male GFP^+/-^;*mdx* mice were injected with AAV vectors, and tissues were harvested and analysed 3 weeks post-injection (Fig. 2b).

**Fig. 2.**
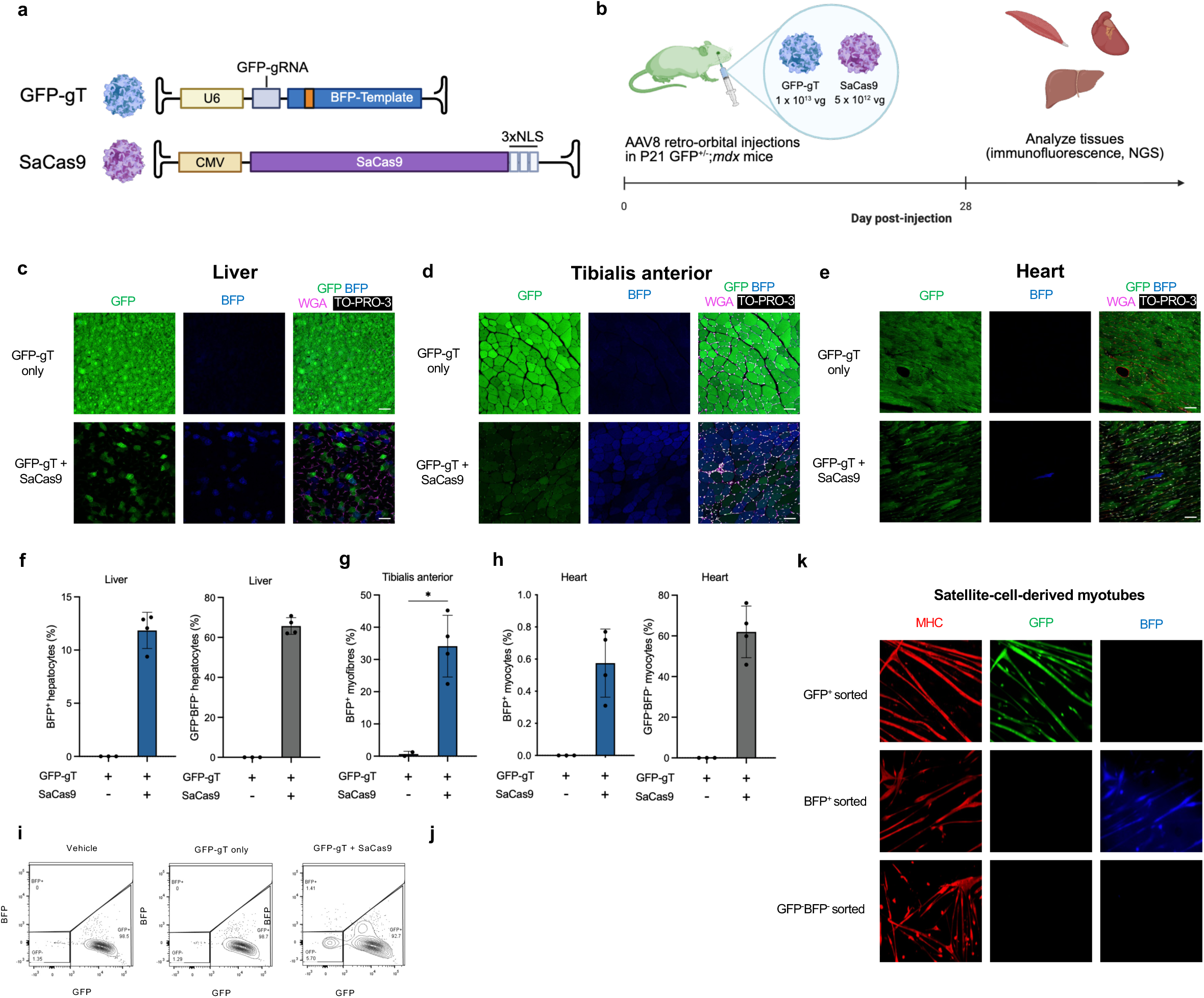
Systemic AAV-GFP-to-BFP enables tracking of *in vivo* CRISPR-NHEJ and CRISPR-HDR in the liver, heart and skeletal muscle of juvenile *mdx* animals. **a,** Adeno**-**associated virus constructs used for in vivo colour switching. **b**, Experimental design. Transgenic P21 GFP^+/-^; *mdx* mice carrying a single CAG-GFP allele were injected with AAV-GFP-gRNA-BFP-Template only (GFP-gT only) or AAV-GFP-gRNA-BFP-Template plus AAV-SaCas9 (GFP-gT + SaCas9). Organs were harvested at day 28 post-injection for immunofluorescence and genomic analyses. **c-f**, Representative immunofluorescence micrographs for detection of CRISPR-NHEJ edited (GFP^-^BFP^-^) and CRISPR-HDR edited (BFP^+^) cells in **b**, liver **d**, heart **e**, and tibialis anterior (TA) muscle after systemic co-injection of GFP-gT and SaCas9 vectors. Scale bars, 50 µm. Green, GFP; Blue, BFP; Magenta, Wheat germ agglutinin (WGA); White, TO-PRO-3. **f-h**, Quantification of BFP^+^ or GFP^-^BFP^-^ cells in representative micrographs of **c**, liver **d**, heart, or **e,** TA muscle. For liver and heart quantification, three fields per tissue per mouse were assessed. For TA muscle, whole cryosections were segmented and the mean fluorescence intensity of each myofibre was measured. A positivity threshold was applied based on the 99.5^th^ percentile of fibre mean fluorescence intensity in negative controls. BFP^+^ myofibres were quantified from 2,771 segmented myofibres across GFP-gT injected mice, and 6,659 fibres across GFP-gT + SaCas9 mice. The frequency of GFP^-^ myofibres were not quantified in skeletal muscle myofibres due to the high degree of multinucleation in this tissue, which prevents detection of complete loss of GFP fluorescence unless all myonuclei contributing to a myofibre are edited. **i**, Representative flow cytometric analyses of skeletal muscle satellite cells isolated from juvenile *mdx* mice injected retro-orbitally with GFP-gT alone or GFP-gT and SaCas9. **j**, Frequency (%) of in vivo-edited BFP^+^ satellite cells and GFP^-^BFP^-^ satellite cells. **k**, Representative micrographs of myotubes differentiated from FACS-sorted satellite cells isolated from in vivo AAV-injected GFP^+/-^; *mdx* mice. Scale bar, 100 µm. Red, myosin heavy chain (MHC); Green, GFP; Blue, BFP. GFP-gT only, n = 3; GFP-gT + SaCas9, n = 4. Data are presented as mean ± SD. **g** was analysed with the Mann-Whitney *U* test; **j** was analysed with one-way ANOVA with Tukey’s multiple comparisons test; n.s., not significant. \**P* < .05.

Control mice received AAV-GFP-gT alone, while experimental mice received AAV-GFP-gT and AAV-SaCas9. We detected widespread loss of GFP signal and gain of BFP signal in the livers of all experimental mice injected with AAV-GFP-gT and AAV-SaCas9, but not in animals receiving AAV-GFP-gT alone (Fig. 2b). We also generated AAV-GFP-g, a cutting-only control vector encoding the GFP-targeting gRNA but lacking the BFP template (Fig. S3a). After systemic administration of AAV-GFP-g and AAV-SaCas9 (Fig. S3b), BFP^+^ hepatocytes were still absent, indicating that indel-mediated editing does not convert GFP to BFP (Fig. S3c, d). In experimental mice, on average, 65.7 ± 4.2% of hepatocytes were GFP^-^BFP^-^, while 11.9 ± 1.7% of hepatocytes were BFP^+^ (Fig. 2c), consistent with previously reported CRISPR-HDR editing rates in neonatal liver^12,13^. The reciprocal loss of GFP in BFP^+^ hepatocytes supports the specificity of reporter conversion and image-based quantification. Amplicon deep sequencing further confirmed concomitant NHEJ and HDR events in the livers of experimental mice (Fig. S4a, b). Because high-dose systemic AAV delivery has been associated with hepatotoxicity in preclinical and clinical studies for muscular dystrophy^14–17^, we analysed liver cryosections, which did not reveal evidence of hepatotoxicity (Fig. S5). These studies confirm the sensitivity and accuracy of our fluorescence imaging-based system for quantifying CRISPR editing events in vivo without the need for immunostaining or signal amplification in sectioned tissues.

Skeletal muscle is largely post-mitotic, composed primarily of multinucleated myofibres formed by fusion of myogenic precursors derived from satellite cells. We and others have utilized AAV-CRISPR-mediated NHEJ in muscle to rescue the *Dmd* reading frame and recover dystrophin expression and function in dystrophic *mdx* mice by deleting or skipping *Dmd* exon 23^18–20^. However, prior attempts at AAV-CRISPR-mediated HDR in muscle^21^ produced negligible editing (∼0.18% alleles edited) when using regulatory regions optimized for expression in post-mitotic fibres (CK8). We therefore evaluated CRISPR-HDR in GFP^+/-^;*mdx* skeletal muscle following systemic dual delivery of AAV-GFP-gT and AAV-SaCas9, under the control of broadly active regulatory elements with high expression in muscle fibres and their precursors (Fig. 2a). Strikingly, we observed widespread BFP^+^ myofibres in the tibialis anterior (TA) muscles of all experimental mice (Fig. 2d, S6a). In contrast, BFP^+^ myofibres were absent in AAV-GFP-gT only (Fig. 2d, S6a) and cutting-only (Fig. S3e, f) controls. Across all experimental mice, 34.1 ± 9.6% of myofibres were BFP^+^, indicating robust presence of HDR-edited genomes (Fig. 2g). As expected, GFP^-^ myofibres were rarely detected, since each myofibre contains hundreds of myonuclei, the majority of which would need to be edited to abolish GFP expression. Notably, a small proportion of HDR-edited nuclei within multinucleated myofibres appears sufficient to drive expression of edited genomes, since fluorescence-based BFP counts were ∼35-fold higher than sequencing-based HDR estimates in TA muscle (Fig. 2g, S3f).

The high percentage of BFP^+^ muscle fibres detected in our study led us to ask whether our system might target endogenous skeletal muscle stem cells (satellite cells), whose progeny could subsequently incorporate into myofibres and amplify the editing effect. We isolated muscle stem cells from AAV-GFP-to-BFP injected mice using a well-established cell surface marker profile (Ter119^-^ CD45^-^ Mac1^-^ Sca1^-^ CXCR4^+^ β1-integrin^+^)^9,10,22^. Consistent with previous data from our group^18^, approximately 5% of FACS-isolated muscle stem cells were GFP^-^BFP^-^, consistent with in vivo GFP disruption by NHEJ (Fig. 2i, j). Importantly, we also detected a small BFP^+^ population (0.84 ± 0.5%), indicative of in vivo HDR editing (Fig. 2i, j). The BFP^+^GFP^-^phenotype in this population was validated by re-sorting culture-expanded cells (Fig. S7) and by deep sequencing (Fig. S4a, b). To test the myogenic function of these in vivo edited satellite cells, they were expanded ex vivo and induced to differentiate. Both in vivo NHEJ- and HDR-edited satellite cells retained the capacity to fuse to form GFP^-^BFP^-^ and BFP^+^ myotubes, respectively (Fig. 2k).

To test whether the highly proliferative environment of *mdx* skeletal muscle might impact rates of HDR editing in vivo, we next compared colour-switching outcomes in GFP^+/-^;*mdx* mice with outcomes in GFP^+/-^ wild-type (GFP^+/-^;WT) controls. In contrast to GFP^+/-^;*mdx* mice, myofibres with BFP fluorescence were not detected in GFP^+/-^;WT mice injected with both AAV-GFP-gT and AAV-SaCas9 (Fig. S8c-e). These data suggest that HDR-mediated editing is significantly enhanced by the microenvironment of dystrophic muscle.

Beyond skeletal muscle, cardiac muscle represents a major therapeutic target for in vivo gene correction, as a wide range of inherited diseases compromise myocardial function. However, the postnatal heart exhibits limited proliferative activity and markedly poor regenerative capacity^23,24^. While we and others have documented AAV-CRISPR-mediated in vivo gene disruption in the heart in mice, the efficiency with which CRISPR-induced DSBs in cardiomyocytes can be resolved through HDR versus NHEJ remains unclear^18–21,25^. In P21 mice injected with both AAV vectors, 62.0 ± 12.7% of cardiomyocytes lost GFP signal, indicating high levels of NHEJ-mediated disruption of the genomic GFP sequence (Fig. 2e, h), whereas BFP was present in only 0.58 ± 0.21% of cardiomyocytes (Fig. 2e, h). Neither GFP disruption nor BFP fluorescence were detected in control mice (Fig. 2e, h). These results demonstrate that while NHEJ-mediated gene disruption occurs efficiently in the postnatal heart, HDR-mediated gene correction remains markedly limited in cardiomyocytes at this stage.

Because cellular proliferation is a major determinant of HDR efficiency, we hypothesized that the paucity of proliferating cardiomyocytes after P21 might contribute to the low HDR rates observed in cardiac versus skeletal muscle at this time point^23,24,26^. To determine whether administration of AAVs in younger mice would increase HDR editing efficiency and to assess the impact of underlying dystrophic pathophysiology in *mdx* mice^27^, we administered AAV-GFP-to-BFP vectors intraperitoneally to GFP^+/-^;*mdx* or GFP^+/-^;WT mice at postnatal day 3 (P3) (Fig. S9a, b). Control animals received AAV-GFP-gT alone, whereas experimental mice received AAV-GFP-gT together with AAV-SaCas9. HDR-mediated BFP expression was detected in 9.9 ± 1.7% of hepatocytes in the GFP^+/-^;*mdx* experimental group and 9.4 ± 4.3% in the GFP^+/-^;WT experimental group, comparable to values observed after P21 injections (Fig. S9c, d). However, the fraction of NHEJ-modified GFP^-^BFP^-^ hepatocytes was lower in neonatally-injected mice (Fig. S9c, d) than in P21 mice (Fig. 2c, f), with frequencies of 23.1 ± 0.7% GFP^+/-^;*mdx* experimental group and 31.8 ± 9.1% in the GFP^+/-^;WT experimental group. These differences may reflect the higher proliferative activity of early neonatal hepatocytes, which would cause more rapid dilutional loss of the non-integrating AAV episomes in neonatal animals^28^.

We next evaluated HDR rates in cardiac and skeletal muscles after systemic administration of AAV-GFP-to-BFP to P3 neonates (Fig. S9a-b). In the heart, HDR-generated BFP^+^ cells were detected at higher frequencies after P3 versus P21 injection, comprising 2.8 ± 1.7% and 3.9 ± 0.6% of cardiomyocytes in GFP^+/-^;*mdx* and GFP^+/-^;WT experimental mice, respectively (Fig. S9e, f and Fig. 2h). The frequency of NHEJ-derived GFP^-^BFP^-^ cells was similar in P3- and P21-injected mice, accounting for 62.4 ± 12.0% of quantified cardiomyocytes in GFP^+/-^;*mdx* mice and 66.9 ± 4.4% in GFP^+/-^;WT mice (Fig. S9e, f). These data suggest that the age-dependent differences in cardiac HDR editing rates are unlikely to be explained by differences in AAV transduction efficiency. In contrast to the heart, skeletal muscle showed no detectable gain of BFP in myofibres in either GFP^+/-^;*mdx* or GFP^+/-^;WT mice administered AAV-GFP-to-BFP (Fig. S9g, h), consistent with the rarity of BFP^+^ skeletal muscle satellite cells in these same muscles (Fig. S10a, b). Loss of GFP could not be evaluated due to the confounding effect of myofibre multi-nucleation, as discussed above. Together, these data reveal discrete, developmentally-timed restrictions on the capacity for in vivo HDR-mediated repair in striated muscle, suggesting that the timing of AAV-CRISPR injection can bias editing toward or away from specific tissues.

Finally, to examine the therapeutic potential of AAV-mediated CRISPR-HDR, we investigated whether HDR could be applied to precisely correct the nonsense mutation in *Dmd* exon 23 in *mdx* mice^29^, which model human Duchenne muscular dystrophy (DMD) and exhibit muscle-wasting caused by the loss of dystrophin protein. We designed a SauriCas9^30^-compatible sgRNA targeting the nonsense mutation and designed a promoterless *Dmd* template carrying the corrective 2989T>C substitution, four synonymous wobble substitutions, and symmetric 900 base pair homology arms (Fig. 3a). Symmetric homology arm design has been shown to be optimal for HDR^31^, and the presence of wobble sequences in the edited DNA both discriminates the template-derived sequences from the endogenous *mdx* genome and disrupts gRNA recognition after HDR editing^32^. AAVs were generated using this *mdx*-gRNA-*Dmd*-Template construct or SauriCas9 and packaged with AAV serotype 8. *Dmd-*CRISPR-HDR AAVs were injected retro-orbitally into juvenile (P21) male *mdx* mice (Fig. 3b), with control mice receiving either AAV-*mdx*-gRNA-*Dmd*-Template alone (AAV-*Dmd*-gT) or AAV-SauriCas9 alone, and experimental mice receiving AAV-*Dmd-*gT plus AAV-SauriCas9 (Fig. 3a).

**Fig. 3.**
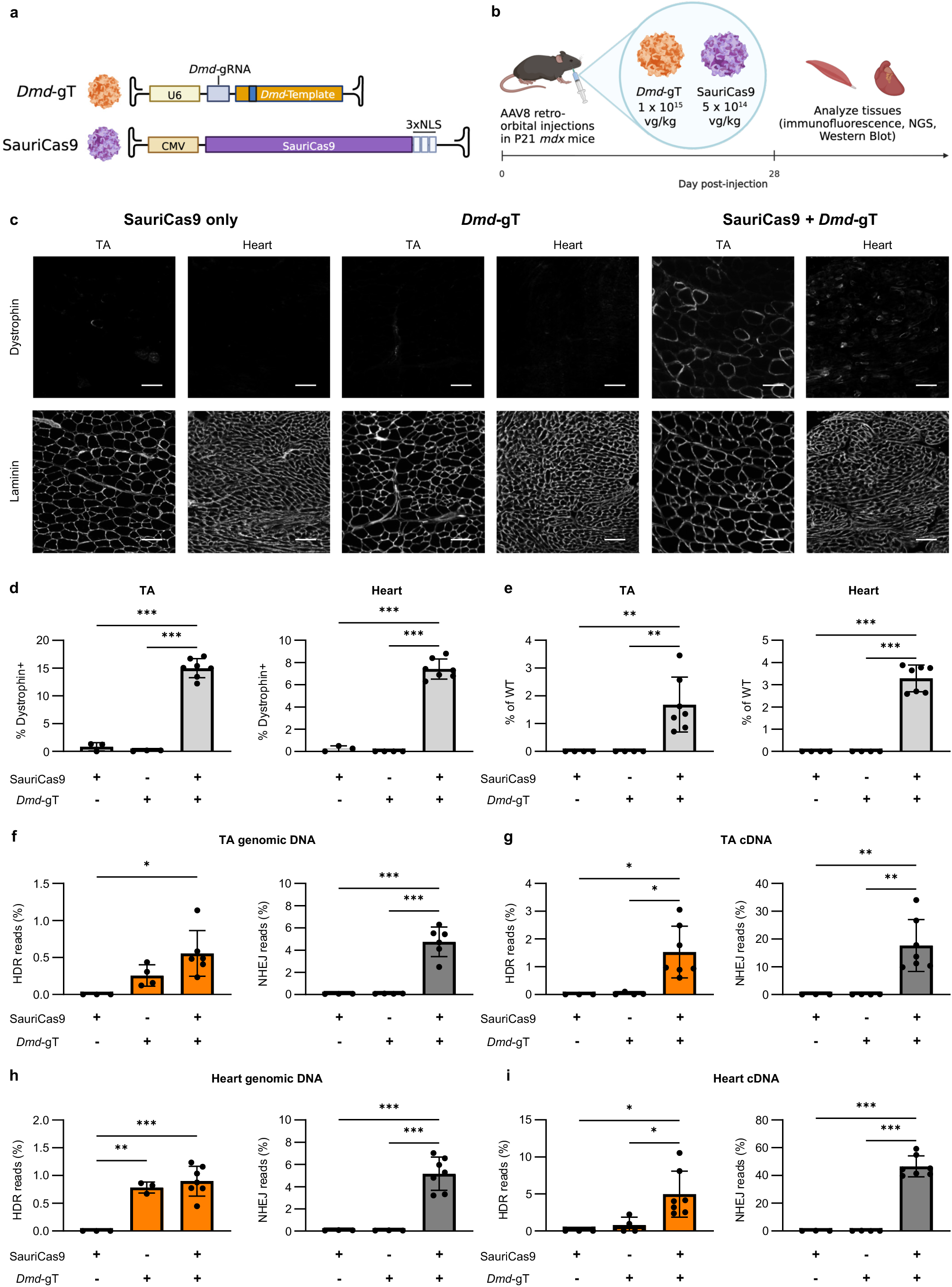
Systemic dual delivery of AAV-SauriCas9 and AAV-*mdx*-gRNA-*Dmd*-Template efficiently restores dystrophin expression in *mdx* mice. **a,** Adeno-associated virus constructs used for in vivo *mdx* mutation correction. **b**, Experimental design. Juvenile (P21) male *mdx* mice were injected with AAV-*mdx*-gRNA-*Dmd*-Template only (*Dmd*-gT; 1 x 10^15^ vg/kg), AAV-SauriCas9 only (SauriCas9; 5 x 10^14^ vg/kg), or AAV-*mdx*-gRNA-*Dmd*-Template plus AAV-SauriCas9 (*Dmd*-gT (1 x 10^15^ vg/kg) + SauriCas9 (5 x 10^14^ vg/kg)). Organs were harvested at day 28 post-injection for immunofluorescence and genomic analyses. **c**, Representative immunofluorescence micrographs for detection of dystrophin (top) and laminin (bottom) expression in TA and heart after systemic co-injection of *Dmd*-gT and SauriCas9 vectors or either vector alone. Representative single-channel images from the same field are displayed for each tissue and condition, as indicated. Scale bar, 100 µm. **d**, Quantification of % dystrophin-positive myofibres or cardiomyocytes in representative micrographs, **c**, calculated by dividing the number of dystrophin-positive cells by the number of laminin-positive cells (total cell number). **e**, Semi-quantitative analysis of dystrophin protein expression in TA and heart of *mdx* mice after systemic co-injection of AAV-*Dmd*-gT and AAV-SauriCas9 vectors or either vector alone. Representative full blots are shown in Fig. S12. **f-i**, Genomic and cDNA from TA and heart were used for amplicon sequencing at the *mdx* mutation site. Representative allele frequency tables are shown in Fig. S14. The frequency (%) of reads with HDR or insertion/deletion (indel) edits in TA at genomic DNA (**f**) and cDNA (**g**) levels and heart at genomic DNA (**h**) and cDNA (**i**) levels. SauriCas9 only, n = 3; *Dmd*-gT, only, n = 3. SauriCas9 *Dmd*-gT, n = 6 in d and 7 in e-i, data are presented as mean ± SD. **d, f, g, h, i,** and **j** are analyzed with one-way ANOVA. *, **, and *** indicate *P* < .05, < .01, and < .001.

Consistent with results from the colour-switching system (Fig. 2), experimental mice showed widespread recoding of the targeted genomic locus, with dystrophin^+^ myofibres and cardiomyocytes detected in the TA muscles and hearts of all experimental mice (Fig. 3c). IHC analysis of Dystrophin-Associated Protein Complex (DAPC) components further confirmed reconstitution in dystrophin^+^ myofibres (Fig. S11). In contrast, no dystrophin^+^ myofibres or cardiomyocytes were detected in controls (Fig. 3c-d). On average, 15.0 ± 1.7% and 7.4 ± 0.9% of myofibres and cardiomyocytes, respectively, were dystrophin^+^ in experimental mice (Fig. 3d).

Western blot analysis also showed that dystrophin protein levels in the TA and heart of experimental mice were 1.7 ± 1.0% and 3.3 ± 0.6% of wild-type mice, respectively (Fig. 3e).

To quantify editing outcomes, genomic DNA and mRNA from the TAs and hearts of control and experimental mice were analyzed by amplicon sequencing of the *mdx* locus (Fig. S13)., we applied a similar strategy for cDNA analysis. Using amplicon sequencing, we determined that 0.6 ± 0.3% and 4.8 ± 1.3% of TA genomic sequences, and 1.5 ± 0.9% and 17.0 ± 9.3% of TA transcripts, were HDR- and NHEJ-edited, respectively (Fig. 3f-g). In heart, 0.9 ± 0.3% and 5.2 ± 1.5% of genomic sequences and 5.0 ± 3.1% and 46.6 ± 7.5% of transcript sequences were HDR-and NHEJ-edited, respectively (Fig. 3h-i). For muscle stem cells, we detected HDR-edited transcripts in 5 out of 7 experimental mice (Fig. S15), providing evidence that therapeutic HDR can occur in this key muscle progenitor population.

Having demonstrated recovery of intact dystrophin sequence by CRISPR-HDR, we next sought to determine whether delivery via the engineered muscle-tropic MyoAAV vector^34^ might offer benefits for reducing the viral dose required, mitigating risk of therapy-associated liver toxicity or other adverse effects associated with the high-dose AAV delivery. Such dose-dependent toxicities have constrained recent DMD gene therapy trials^17,35,36^. Notably, MyoAAV vectors show preserved therapeutic efficacy with reduced systemic doses in other gene therapy and gene editing applications^34^, and demonstrate enhanced transduction of endogenous muscle satellite cells in juvenile- (P21) injected mice (Fig. S16). We therefore evaluated MyoAAV-mediated CRISPR-HDR in P21 *mdx* mice by injection of AAV-*mdx*-gRNA-*Dmd*-Template and AAV-SauriCas9 packaged in MyoAAV2A, the highest-performing MyoAAV subtype for systemic muscle delivery in mice^37^. Experimental mice were injected with both MyoAAV2A-*mdx*-gRNA-*Dmd*-Template and MyoAAV2A-SauriCas9, while control P21 *mdx* mice received either MyoAAV2A-*mdx*-gRNA-*Dmd*-Template or MyoAAV2A-SauriCas9 alone (Fig. 4a-b). Notably, compared to AAV8-mediated *Dmd* HDR correction (Fig. 3), MyoAAV-mediated HDR studies were performed at a 25-fold lower systemic dose. Four weeks after injection, widespread dystrophin^+^ myofibres and cardiomyocytes were observed in TA muscles and hearts, respectively, of experimental, but not control, mice (Fig. 4c). On average, 22.8 ± 5.0% of myofibres and 11.8 ± 1.8% of cardiomyocytes were dystrophin^+^ in experimental mice injected with Dmd CRISPR-HDR MyoAAVs (Fig. 4d). Analysis of DAPC components further confirmed reconstitution in dystrophin^+^ myofibres (Fig. S11). Western blotting indicated that dystrophin expression levels in the TA and heart of experimental mice were 0.5 ± 0.4% and 0.6 ± 0.4% of wild-type mice (Fig. 4e). Amplicon sequencing revealed that 0.2 ± 0.1% and 4.3 ± 1.1% of genomic sequences, and 0.7 ± 0.3% and 15.3 ± 3.7% of transcripts, were HDR- and NHEJ edited, respectively, in TA muscles (Fig. 4f-g). In heart, 0.2 ± 0.1% and 4.1 ± 1.0% of genomic sequences and 1.6 ± 0.9% and 16.0 ± 4.2% of transcripts were HDR- and NHEJ-edited, respectively (Fig. 4h-i). HDR-edited transcripts were also detected in satellite cells in 3 out of 8 experimental mice (Fig. S15). Together, these results demonstrate full recovery of dystrophin coding sequence using MyoAAV at a 25-fold lower viral dose compared to AAV8.

**Fig. 4.**
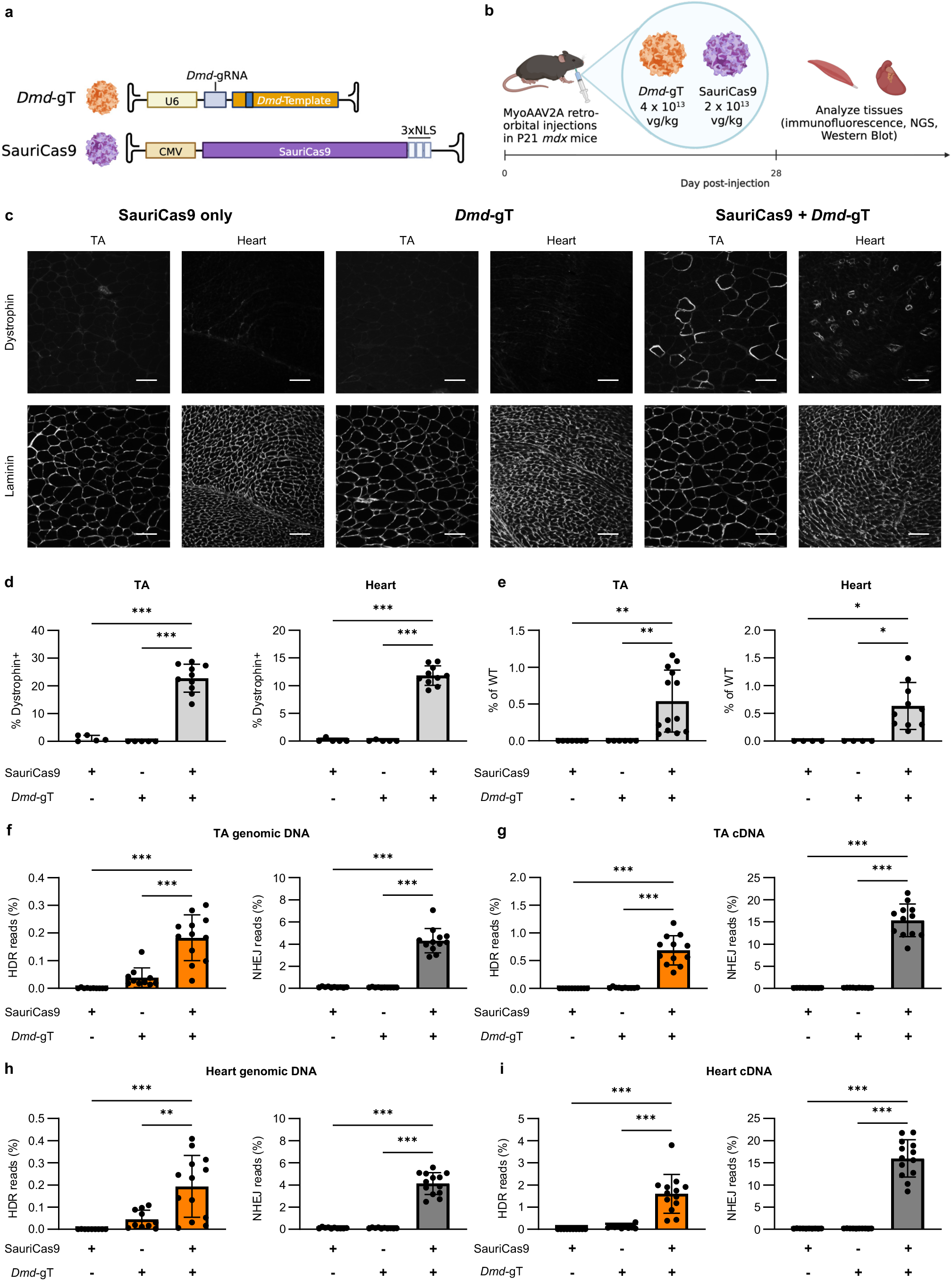
Systemic dual delivery of MyoAAV-SauriCas9 and MyoAAV-*mdx*-gRNA-*Dmd*-Template efficiently restores dystrophin expression in *mdx* mice. **a,** Adeno-associated virus constructs used for in vivo *mdx* mutation correction. **b**, Experimental design. Juvenile (P21) male *mdx* mice were injected with MyoAAV-*mdx*-gRNA-*Dmd*-Template (*Dmd*-gT; 4 x 10^13^ vg/kg), MyoAAV-SauriCas9 (SauriCas9; 2 x 10^13^ vg/kg), or MyoAAV-*mdx*-gRNA-*Dmd*-Template plus MyoAAV-SauriCas9 (*Dmd*-gT (4 x 10^13^ vg/kg) + SauriCas9 (2 x 10^13^ vg/kg)). Organs were harvested at day 28 post-injection for immunofluorescence and genomic analyses. **c**, Representative immunofluorescence micrographs for detection of dystrophin (top) and laminin (bottom) expression in TA and heart after systemic co-injection of *Dmd*-gT and SauriCas9 vectors or either vector alone. Representative single-channel images from the same field are displayed for each tissue and condition, as indicated. Scale bar, 100 µm. **d**, Quantification of % dystrophin-positive myofibres or cardiomyocytes in representative micrographs, **c**, calculated by dividing the number of dystrophin-positive cells by the number of laminin-positive cells (total cell number). **e**, Semi-quantitative analysis of dystrophin protein expression in TA and heart of *mdx* mice after systemic co-injection of MyoAAV-*Dmd*-gT and MyoAAV-SauriCas9 vectors or either vector alone. Representative full blots are shown in Fig. SX. (**f-i**) Genomic and cDNA from TA and heart were used for amplicon sequencing at the *mdx* mutation site. Representative allele frequency tables are shown in Fig. SX. The frequency (%) of reads with HDR and insertion/deletion (indel) edits in TA at genomic DNA (**f**) and cDNA (**g**) levels and heart at genomic DNA (**h**) and cDNA (**i**) levels. SauriCas9 only, n = 5 in d, 7 in e TA and 4 in e heart, and 10 in f-i; *Dmd*-gT, n = 5 in d, 6 in e TA and 4 in e heart, and 10 in f-i; SauriCas9 + *Dmd-*gT, n = 10 in d and e heart, 13 in e TA, 12 in f-g, and 13 in h-i; *Dmd*-gT, only, n = 3. SauriCas9 *Dmd*-gT, n = 6 in d and 7 in e-i. Data are presented as mean ± SD. d, f, g, h, i, and j are analyzed with one-way ANOVA. *, **, and *** indicate *P* < .05, < .01, and < .001.

In conclusion, our study establishes a simple fluorescent-based platform to track NHEJ and HDR gene editing outcomes in vivo across a variety of tissues, and reveals an unexpected opportunity for precise, targeted gene replacement by HDR in skeletal and cardiac muscles, both largely post-mitotic tissues that have been widely considered to be inaccessible by this approach^21,38^. Whether similar developmentally controlled windows of CRISPR-HDR accessibility exist for other cell types will be an important question for future investigation. Our study also provides the first demonstration of successful HDR-editing in tissue stem cells within their native niche, which will uniquely enable directed manipulation of stem cell genomes therapeutically and experimentally, without the need to isolate, expand or transplant these rare cells. We also extend our findings to achieve therapeutic correction of disease-relevant gene loci, with restoration of intact dystrophin coding sequence in skeletal and cardiac muscle, with muscle-targeted CRISPR-HDR delivery via MyoAAV enabling a 25-fold reduction in the systemic AAV dose, compared to AAV8. Ultimately, the ability to inscribe enduring, precise genome modification in the neonatal mammalian heart and postnatal mammalian skeletal muscle satellite cells opens exciting new avenues for future therapeutic interventions for many currently intractable cardiac and muscle diseases.

## Author Contributions

B.L.P., K.H.L., A.L., K.Z., and A.J.W. conceived the study and designed experiments. K.Z. performed in vitro and ex vivo reporter validation. B.L.P. and K.Z. performed molecular cloning and vector construction. B.L.P., K.H.L., A.L., K.Z., C.L.R., J.G., K.M., H.K., N.H., A.D.B., T.L., M.M., and R.E. conducted animal experiments and tissue harvests. K.Z. and B.L.P. carried out functional myogenic assays. B.L.P., K.H.L., A.L., C.L.R., A.D.B., and T.L. performed cryosectioning, image acquisition, and quantification. B.L.P., K.H.L., A.L., M.M., R.E., K.M., and K.Z. performed satellite cell isolation, fluorescence-activated sorting, and culture. K.M. carried out histological analysis. R.E. performed Western blot analyses. B.L.P., A.L., and M.M. prepared amplicons for deep sequencing. M.F. and K.H.L. performed analysis of the deep amplicon sequencing results. S.A. generated reagents and assisted with in vitro experiments. R.X., L.V., U.S.P., K.H.L., A.L., and M.K.C. generated and titered AAVs. A.J.W. supervised the study. B.P., K.H.L., A.L., and A.J.W. wrote the manuscript with input from all co-authors.

## Acknowledgements

We thank J. LaVecchio and S. Ionescu at the HSCI/HSCRB Flow Cytometry Core for assistance with fluorescence-activated cell sorting; the SERI/MEEI Gene Transfer Vector Core and Gene Therapy Program Preclinical Vector Core at the University of Pennsylvania for generating AAVs; A. Almada and M. Tabebordbar for discussions and critically reading the manuscript; C. Macgillivray at the HSCRB Histology Core for assistance with histology; the Harvard Center for Biological Imaging for microscopy infrastructure and support; the MGH CCIB DNA Core for assistance with sequencing; and Wagers lab members for support and suggestions. This work was funded in part by grants from Harvard University (Dean’s Fund and Star Family Challenge Award), the New York Stem Cell Foundation, and the Glenn Foundation for Medical Research (to A.J.W.).

## METHODS

No statistical methods were used to predetermine sample size. Mice used for in vivo experiments were randomized for injections of AAVs. Investigators responsible for cryosectioning, image acquisition, quantification, and sequencing were blinded to sample allocation.

### Animals

Hemizygous GFP transgenic mice, carrying a single transgenic allele, were generated by crossing CAG-GFP mice^5^ with either C57BL/6J or C57BL/10ScSn-*Dmd^mdx^*/J (*mdx*) (Jackson Labs). C57BL/10ScSn-*Dmd^mdx^*/J (JAX stock #001801) and C57BL/10ScSnJ (JAX stock #000476) mice were purchased from the Jackson Laboratory. Postnatal day 3 (P3) GFP*^+/-^*;*mdx* and GFP*^+/-^*;WT pups (both male and female) were used for neonatal intraperitoneal (IP) injections and 3-week-old male GFP*^+/-^*;*mdx* mice were used for juvenile intravenous (retro-orbital) injections for GFP-to-BFP colour switching experiments. 3-week-old male *mdx* mice were used for juvenile intravenous (retro-orbital) injections for *mdx* mutation correction experiments. Mice were maintained at the Harvard Biological Research Infrastructure according to animal care and experimental protocols approved by the Harvard University Institutional Animal Care and Use Committee (IACUC).

### AAV production and administration

For AAV8 used for GFP-to-BFP colour switching experiments in Fig. 2 and Fig. S3-S6, and S9-S10, vectors were produced and titered by the Gene Transfer Vector Core (GTVC) at the Grousbeck Gene Therapy Center at the Schepens Eye Research Institute and Massachusetts Eye and Ear Infirmary (SERI/MEEI). For AAV8 used for GFP-to-BFP colour switching experiments in Fig. S7-S8 and *mdx* mutation correction experiments in Fig. 3 and S11-15, vectors were produced and titered by Gene Therapy Program Preclinical Vector Core at the University of Pennsylvania. AAV were packaged as previously described. Briefly, semi-confluent HEK293 cells were transfected with rep2-cap8 packaging construct, an adenoviral helper function plasmid, and the ITR flanked transgene construct. Three days following transfection, media and cells were harvested, underwent lysis and benzonase digestion for removal of non-particle associated DNA. Particles were purified and concentrated using tangential flow filtration, iodixanol density centrifugation, and buffer exchange into a PBS-based buffer solution. For MyoAAV used for *mdx* mutation correction experiments in Fig. 4, S11, and S16-18 vectors were produced in-house as previously described^39^. In contrast to outsourced AAV productions, in-house produced particles were purified and concentrated using POROS™ GoPure™ AAVX Pre-packed Column (Thermo Scientific) and high-performance lipid chromatography (HPLC).

For neonatal (P3) intraperitoneal AAV8 injections, control mice received 3 x 10^12^ viral genome (vg) of AAV-GFP-gRNA-BFP-template alone, while experimental mice received 3 x 10^12^ vg of AAV-GFP-gRNA-BFP-template plus 1 x 10^12^ vg of AAV-SaCas9. Virus was diluted in 75mL of vehicle (PBS with 35mM NaCl) for each injection. Mice were analyzed 28 days post-injection.

For juvenile (P21) retro-orbital AAV8 injections used for GFP-to-BFP colour switching, gRNA-BFP-Template only control mice received 1 x 10^13^ vg of AAV-GFP-gRNA-BFP-template alone, and experimental mice received 1 x 10^13^ vg of AAV-GFP-gRNA-BFP-template plus 5 x 10^12^ vg of AAV-SaCas9. For repeat juvenile (P21) retro-orbital AAV8 injections used for assessing whether Cas9 cleavage without an HDR donor template in vivo can result in BFP, cutting-only control mice received 1 x 10^13^ vg of AAV-GFP-gRNA plus 5 x 10^12^ vg of AAV-SaCas9. For juvenile (P21) retro-orbital AAV8 injections used for *mdx* mutation correction experiments, control mice either received 1 x 10^15^ vg/kg of AAV-*mdx*-gRNA-*Dmd*-template alone or 5 x 10^14^ vg/kg of AAV-SauriCas9 alone, and experimental mice received 1 x 10^15^ vg/kg of AAV-*mdx*-gRNA-*Dmd*-template plus 5 x 10^14^ vg/kg of AAV-SauriCas9. For juvenile (P21) retro-orbital MyoAAV injections used for mdx mutation correction experiments, control mice either received 4 x 10^13^ vg/kg of MyoAAV-*mdx*-gRNA-*Dmd*-template alone or 2 x 10^13^ vg/kg of MyoAAV-SauriCas9 alone, and experimental mice received 4 x 10^13^ vg/kg of MyoAAV-*mdx*-gRNA-*Dmd*-template plus 2 x 10^13^ vg/kg of MyoAAV-SauriCas9. Virus was diluted in vehicle (PBS with 35mM NaCl and 0.001% Pluronic F-68) for a maximum volume of 100 µL for each injection. Mice were analyzed 28 days post-injection.

### Gene editing constructs

The AAV-SaCas9 plasmid was previously described^18^. AAV-GFP-gRNA-BFP-Template plasmid was generated by Gibson assembly of the pZac2.1 AAV vector with three inserts. The vector was double digested by HindIII-HF and NotI-HF (NEB). Insert fragment 1 (U6-GFP-gRNA) was PCR amplified from a plasmid containing U6-GFP-gRNA. Insert fragment 2 (BFP) was PCR-amplified from a BFP sequence synthesized as a gBlock (IDT). Insert fragment 3 (polyA) was PCR-amplified from genomic DNA of the CAG-GFP transgenic animal. Two base substitutions on the BFP template enable the colour switch (from green to blue fluorescence), block the seed sequence of the protospacer to reduce further Cas9-mediated cleavage, and generate a restriction fragment length polymorphism (RFLP) detectable by BtgI restriction enzyme digestion.

### Satellite cell isolation, culture, and differentiation

Satellite cells for *ex vivo* gene editing were isolated as previously described^9^. For isolation of in vivo edited satellite cells, triceps, abdominal and hindlimb muscles from half of the body were harvested and minced using scissors, then subjected to two rounds of digestion with 0.2% Collagenase type II and 0.05% Dispase in DMEM (GIBCO) at 37°C (for 15 min, then 10 min). Enzymes were inactivated by addition of FBS, and cells were centrifuged and filtered through 70 µm strainers before staining for 30 min with an antibody cocktail containing APC-Cy7-CD45 (Biolegend, 1:200), APC-Cy7-CD11b (Biolegend, 1:200), APC-Cy7-TER119

(Biolegend, 1:200), APC-Sca1 (Biolegend, 1:200), PE-CD29 (Biolegend, 1:100) and Biotin-CD184 (BD Biosciences, 1:100). After primary antibody incubation, cells were washed with staining media (SM, Hank’s Balanced Salt Solution + 2% serum) and then stained for an additional 20 min with Streptavidin PE-Cy7 (Biolegend, 1:200). Finally, cells were washed twice in SM and resuspended in SM containing propidium iodide (PI) to mark dead cells. Satellite cells were sorted using a FACS Aria II (BD Biosciences) based on their lack of PI incorporation and CD45, Ter119, Sca1 and CD11b expression and positive expression of CXCR4 (CD184) and β1-integrin (CD29), a surface marker profile that has been extensively validated^9,10,22^ to select for Pax7^+^ cells with robust myogenic capacity. Separately sorted GFP^+^, BFP^+^, and GFP^-^BFP^-^satellite cells were expanded on collagen type I (1 µg/mL, Sigma) and laminin (10 µg/mL, Invitrogen) coated plates in Growth Media (F10, 20% horse serum, 1% Pen Strep, and 1% Glutamax (Gibco)), supplemented daily with 5 ng/mL bFGF (Sigma). DNA was isolated from subset of the expanded cells was harvested using QuickExtract (Lucigen) and used for genomic PCR and subsequent RFLP and sequencing analysis. Myogenic differentiation was initiated by switching to Differentiation Media (DMEM, 2% horse serum, 1% Pen Strep, 1% Glutamax (Gibco)) for 3-4 days. Cells were fixed by 4% PFA for 20 minutes for imaging.

### Transfection

Satellite cells isolated from male GFP^+/^;*mdx* mice were expanded in culture in Growth Media with daily bFGF supplementation for 2-3 weeks and then re-plated onto 24 well plates coated with collagen (1 µg/mL) and laminin (10 µg/mL) at 20,000 cells per well. Myoblasts were transfected on day 2 using Lipofectamine 3000 (Invitrogen) per manufacturer’s instructions with GFP-gRNA-BFP-Template plasmid alone for control group or GFP-gRNA-BFP-Template and SaCas9 plasmids at a 5:1 ratio the experimental group (3 independent transfections per group). BFP^+^ and GFP^+^ cells were sorted using a FACS Aria II 5 days after transfection and resorted after an additional 2 weeks of expansion in vitro to confirm fluorescence. Re-sorted cells were then used for in vitro differentiation and in vivo transplantation assays. For testing GFP disruption in GFP^+/-^;*mdx* primary myoblasts, cells were transfected with Lipofectamine only (control) or with plasmids encoding SaCas9 and GFP-gRNA-2 (no BFP template) at 1:1 ratio, as described above. For screening of GFP-targeting gRNAs, GFP^+/-^;*mdx* tail tip fibroblasts (TTFs) were transfected with SaCas9 alone (control) or with SaCas9 plus one of the three gRNAs targeting GFP, using Lipofectamine 3000 per manufacturer’s instructions.

### Myoblast transplantation

One day before myoblast transplantation, 25 mL of *Naja mossambica mossambica* cardiotoxin (0.03 mg/mL, Sigma) was injected to the tibialis anterior (TA) muscles of anesthetized male *mdx* recipient mice. 800,000 GFP^+^, 800,000 BFP^+^ myoblasts or vehicle (PBS) alone was injected into the pre-injured TA muscles (n = 4 TA muscles). The injected TA muscles were harvested 5 weeks post-transplantation for cryosectioning and immunofluorescence.

### Genomic PCR and RFLP analysis

Genomic DNA from tissues, satellite cells and expanded myoblasts was extracted using QuickExtract DNA Extraction Solution (Epicentre/Lucigen) or DNeasy Blood & Tissue Kit (Qiagen) per manufacturer’s protocol. 1-2 mL of solutions extracted with QuickExtract was used per 25 mL PCR reaction by Q5 Hot Start High-Fidelity DNA Polymerase (NEB). For GFP-to-BFP colour switching experiments, the Forward primer GTGCTGTCTCATCATTTTGGC (binds upstream of GFP/BFP start site) and Reverse primer TCGTGCTGCTTCATGTGGTC (binds downstream of Cas9 cutting site and colour switching substitutions) were used to amplify the genomic transgene locus, but not template sequence.

For RFLP analysis, PCR products were purified using QIAquick PCR Purification Kit (Qiagen) and digested with BtgI (NEB), or mock digested with water, before gel electrophoresis on E-Gel EX 2% Agarose Gels (Invitrogen).

### RNA extraction and cDNA synthesis

Muscle tissues were flash frozen with liquid nitrogen and satellite cell pellets were frozen in -20℃ freezer. Muscle tissues were lysed with 1 mL of Trizol (ThermoFisher) in gentleMACS^TM^ M tubes (Miltenyi Biotec) using gentleMACS Octo tissue dissociator (Miltenyi Biotec) with program RNA_02. Following tissue lysis, RNA-containing aqueous phase was obtained following manufacturer’s instructions and then subjected to RNeasy MiniPrep Kit (Qiagen) to extract total RNA. For satellite cell pellets, total RNA was extracted directly using RNeasy MiniPrep Kit (Qiagen). 500 ng of the extracted total RNA was digested with ezDNase (ThermoFisher) and used as template for first-strand cDNA synthesis using SuperScript™ IV First-Strand Synthesis System (ThermoFisher) with oligo d(T)_20_ per manufacturer’s instructions.

### Sanger sequencing and deep amplicon sequencing

Purified genomic PCR products were cloned into the TOPO backbone using the Zero Blunt TOPO PCR cloning kit (Invitrogen) and transformed into One Shot TOP10 Chemically Competent *E. coli* (Invitrogen). Ten discrete clones per condition were analyzed by bacterial colony Sanger sequencing, performed by Genewiz in Cambridge, MA. Sequencing traces were aligned to the GFP transgene using Geneious (Dotmatics). For deep amplicon sequencing of the GFP locus, 8 base pair (bp) barcodes were appended to the genomic PCR primers, and 4-5 uniquely barcoded PCR products were pooled and PCR purified using the QIAquick PCR Purification Kit (Qiagen), followed by analysis at the MGH DNA core using their CRISPR Sequencing service. For *mdx* mutation correction experiments, PCR products were not pooled to ensure sufficient sequencing depth. For *Dmd* exon 23 genomic DNA analyses, PCR amplification was performed with the forward primer targeting a region that is not on the *Dmd* HDR donor template to prevent amplifying AAV genomes containing the *Dmd* HDR donor template. PCR products were gel extracted with QIAquick Gel Extraction Kit (Qiagen) and used as DNA template to amplify amplicons for NGS. For *Dmd* exon 23 cDNA analysis, we performed PCR amplification with the forward primer targeting *Dmd* exon 22 (not on DMD HDR template. Primer sequences are available upon request. For all deep amplicon sequencing samples, PCR products were purified before analysis at the MGH DNA core by CRISPR Sequencing. For deep amplicon sequencing of the GFP locus, each sample yielded between 500 and 10,000 amplicon-aligned reads. BBTools was first used to merge paired R1 and R2 reads into a single contiguous read containing both 5’ and 3’ barcodes. Cutadapt was then used to demultiplex the data based on each sample’s unique barcode. Samples with fewer than 1,000 reads were excluded to ensure sufficient sequencing depth, as previously described for pooled analyses^40^. Deep sequencing results were then analyzed using CRISPResso2. The frequency of indel was determined using the CRISPresso2 output and HDR was determined by manually quantified the reads bearing template sequence (including wobble sequences). Variants with mdx mutation sequence replaced by DMD template sequence (including wobble sequences) as HDR-edited, and all the other variants as non-HDR mismatch.

### Sectioning and immunofluorescent imaging

For GFP-to-BFP colour switching experiments, tissues were dissected and immediately fixed in 4% PFA for 90 minutes at room temperature and then washed with PBS and transferred to 30% sucrose for overnight incubation at 4°C. Submersed tissues were then embedded in O.C.T. compound (Tissue-Tek) and frozen in isopentane in a liquid nitrogen bath. Tissues were cryosectioned using a CM1860 cryostat (Leica Biosciences) at a thickness of 12 µm and blocked with 10% normal goat serum (NGS; Jackson ImmunoResearch) and 2% bovine serum albumin (BSA; Sigma Aldrich) for 1 hour at room temperature. For laminin staining, tissues were incubated in rabbit polyclonal anti-laminin antibody (1:1000; Sigma Aldrich, L9393) overnight in a humidified chamber, followed by 3 x 5 min DPBS washes. Sections were then incubated with goat anti-rabbit Alexa Fluor 555 for 1 hour at room temperature. For wheat germ agglutinin staining, sections were incubated in Wheat Germ Agglutinin Alexa Fluor 555 conjugate for 1 hour at room temperature, followed by 3 x 5 min DPBS washes. Sections were stained with TO-PRO-3 Iodide (Life Technologies) according to manufacturer’s instructions. Images were captured using a Zeiss LSM 880 inverted confocal microscope (ZEISS, Germany). For in vivo AAV-GFP-to-BFP experiments, numbers of BFP^+^, GFP^-^, and total cells were counted manually using ImageJ. For liver and heart, three representative fields with ∼200-350 cells per 20x field were counted for each tissue. For tibialis anterior muscle, stitched whole-cryosection confocal 20x images were analysed using a semiautomated fluorescence intensity-based pipeline described in “Semi-automated myofibre segmentation and fluorescence-based quantification.”

For *mdx* mutation correction experiments, tissues were directly embedded in optimal cutting temperature compound (OCT, Tissue-Tek) and flash-frozen in liquid-nitrogen-chilled isopentane for cryosectioning. Tissues were cryosectioned using CM1860 cryostat (Leica Biosciences) at a thickness of 12 μm. For laminin and dystrophin immunofluorescence staining,muscle sections were permeabilized with 0.2% Triton X-100 in PBS for 10 minutes, followed by 3 x 5 min DPBS washes. Sections were then blocked with 10% normal goat serum (NGS; Jackson ImmunoResearch) and 2% bovine serum albumin (BSA; SigmaAldrich) for 2 hours at room temperature. Sections were then incubated with mouse monoclonal IgG2b anti-dystrophin antibody (MANDYS8, Sigma-Aldrich; 1:100) and rabbit polyclonal anti-laminin antibody (L9393, Sigma-Aldrich; 1:200) prepared in DPBS containing 5% NGS and 0.01% TWEEN-20 for 1 hour at room temperature, followed by 3 x 5 min DPBS washes. Sections were subsequently incubated with goat anti-mouse IgG2b Alexa Fluor 594 (1:1000, Thermo Scientific) and goat anti-rabbit IgG Alexa Fluor 488 (1:250, Invitrogen) in DPBS with 5% NGS and 0.01% TWEEN-20 for 2 hours at room temperature, followed by 3 × 5-min DPBS washes. Finally, sections were stained with Hoescht (1:1000) for 5 minutes at room temperature followed by 3 x 5 min DPBS washes. Images were captured using a Zeiss LSM 880 confocal inverted microscope (ZEISS, Germany) or an ECHO confocal microscope on wide-field mode (ECHO, San Diego, CA, USA). Laminin-positive and dystrophin^+^ myofibres were quantified manually using ImageJ in a blind analysis. Dystrophin expression is reported as the percentage of dystrophin^+^ myofibres relative to the total number of laminin^+^ myofibres.

For α-sarcoglycan, β-sarcoglycan, β-dystroglycan, and nNOS immunofluorescence staining, serial cross-sections of muscle tissues were permeabilized with 0.2% Triton X-100 in DPBS for 10 minutes, followed by 3 x 5 min DPBS washes. Sections were then blocked with 5% normal goat serum (NGS; Jackson ImmunoResearch), 2% bovine serum albumin (BSA; Sigma-Aldrich), 2% protein concentrate and 1 drop/mL M.O.M. Blocking Reagent (M.O.M. Kit; Vector Laboratories), and 0.1% TWEEN-20 in DPBS for 1 hour at room temperature, followed by 3 x 5 min DPBS washes. Sections were incubated overnight at 4°C with one of the following primary antibodies: mouse monoclonal IgG1 anti-α-sarcoglycan (GeneTex; 1:50), mouse monoclonal IgG1 anti-β-sarcoglycan (Leica; 1:100), mouse monoclonal IgG2a anti-β-dystroglycan (Leica; 1:100), or rabbit polyclonal anti-nNOS (ImmunoStar; 1:1,000). All sections were co-stained with mouse monoclonal IgG2b anti-dystrophin antibody (MANDYS8, Sigma-Aldrich; 1:100).

Primary antibodies were diluted in PBS containing 3% BSA, 8% protein concentrate (M.O.M. Kit; Vector Laboratories), and 0.1% Tween-20. Following primary antibody incubation, sections were washed 4 x 10 min with DPBS. Sections were then incubated for 1 hour at room temperature with the appropriate secondary antibodies: goat anti-mouse IgG1 Alexa Fluor 488 (Thermo Scientific; 1:250), goat anti-rabbit IgG Alexa Fluor 647 (Thermo Scientific; 1:250), or goat anti-mouse IgG2a Alexa Fluor 647 (Thermo Scientific; 1:250), together with goat anti-mouse IgG2b Alexa Fluor 594 (Thermo Scientific; 1:1000). Secondary antibodies were diluted in DPBS containing 3% BSA and 0.1% Tween-20. After secondary antibody incubation, sections were washed 4 x 10 min with DPBS. Finally, sections were stained with Hoescht (1:1000) for 5 minutes at room temperature followed by 3 x 5 min DPBS washes.

### Semi-automated myofibre segmentation and fluorescence-based quantification

To avoid pseudoreplication, for TA muscle, a single representative stitched micrograph of a whole cryosection was assessed per mouse. Immunofluorescence micrographs of whole tibialis anterior cryosections were segmented with Cellpose-SAM v4.1.1^41^. Cellpose-SAM-derived masks were converted to ImageJ region of interest (ROI) files using LabelsToRois^42^ and were manually curated to remove objects that did not represent myofibres, merged myofibres, partial myofibres at micrograph edges and damaged or folded tissue. To exclude sarcolemmal and extracellular autofluorescence, ROIs were eroded by 7 pixels using LabelsToRois. For each image, ten ROIs of background fluorescence were placed in regions without tissue or artifacts.

For each image and channel, the median background intensity across the ten background ROIs was subtracted from the per-myofibre median fluorescence intensity to account for image-specific background signal. Corrected median fluorescence intensity values for every segmented myofibre were used for downstream analyses.

### Western blot analysis

Protein lysates were prepared from frozen tibialis anterior (TA) or heart tissues using a gentleMACS Octo Dissociator (Miltenyi Biotec) with 500 µL of RIPA buffer per tissue sample (Thermo Scientific) along with 1% Halt protease and phosphatase inhibitor cocktail (Thermo Scientific) and 1% EDTA for 53 sec at 2753 rpm using gentleMACS M tubes. Total protein concentration was measured using the Qubit Protein Broad Range Assay Kit (Thermo Scientific) following the manufacturer’s instructions.

For Western blot analysis, equal amounts of total protein were resolved on NUPAGE 3–8% Tris-Acetate precast gels (Thermo Scientific) using the NuPAGE electrophoresis system (Invitrogen PowerEase 350W) at 150 V for approximately 75 min. Proteins were transferred onto 0.2 µm nitrocellulose membranes with a Trans-Blot Turbo Transfer System (Bio-Rad) using Turbo Transfer Packs. The standard high-molecular-weight transfer protocol was modified to a constant current of 2.0–2.5 A for 15 min.

Membranes were then blocked in 5% (w/v) non-fat milk (Bio-Rad) in TBST (1X Tris Buffered Saline (Bio-Rad) with 0.1% TWEEN-20) for 1 hour at room temperature. Membranes were incubated overnight at 4°C with rabbit anti-dystrophin primary antibody (Abcam, ab154168; 1:2000) prepared in 5% milk in TBST. Following thorough washes with TBST, membranes were then incubated for 1 hour at room temperature with an HRP-conjugated anti-rabbit secondary antibody (Thermo Scientific). After additional TBST washes, chemiluminescent detection was performed using SuperSignal West Femto Maximum Sensitivity Substrate (Thermo Scientific), and images were acquired with a ChemiDoc MP Imaging System (Bio-Rad). Following chemiluminescent detection, membranes were incubated with Coomassie blue staining solution (0.1% Coomassie Brilliant Blue G-250 (Thermo Scientific) in 40% methanol and 10% acetic acid) for 10 minutes. Membranes were briefly rinsed with water, dried overnight, and imaged. Band intensities were quantified using ImageJ software (NIH).

### Statistical analysis

GraphPad Prism 11.0.2 software was used for performing statistical analysis. Unpaired two-tailed t test, one-way ANOVA with Tukey’s multiple comparisons test, Mann-Whitney test, and two-tailed Fisher’s exact test were performed. For all tests, a *P* value of less than .05 was considered statistically significant. *P* values and methods can be found in corresponding figure legends.

### Data availability

Raw data are available from the corresponding author on reasonable request.

---

**Fig. S1.**
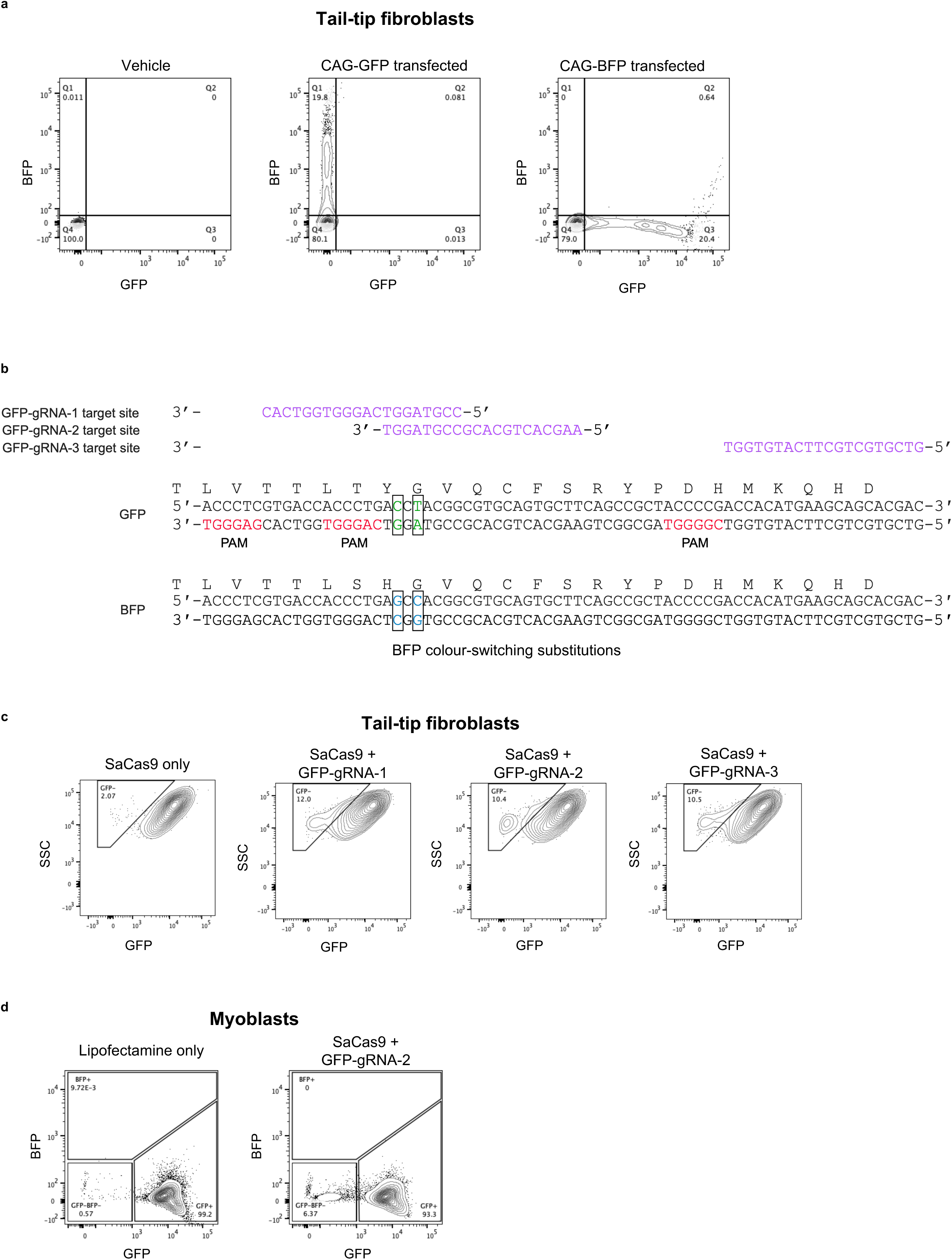
Validation of GFP-to-BFP colour switching reporter system. **a**, Representative flow cytometry plots of tail-tip fibroblasts (TTFs) isolated from *mdx* mice transfected with either CAG-GFP or CAG-BFP at 3 days post-transfection. **b**, Design and sequence of colour switching substitutions and GFP-gRNAs targeted GFP. Two base substitutions cause spectral shift and create a BtgI site for restriction fragment length polymorphism (RFLP) analysis. Three SaCas9-compatible gRNAs targeting GFP near the substitution site were selected. GFP-gRNA-2 cuts closest to the desired colour-determining bases and protospacer recognition by the GFP-gRNA-2 spacer is blocked by HDR substitutions, which protects the BFP template and genomic HDR product from further Cas9-induced double-stranded breaks. **c**, Representative flow cytometry plots of GFP^+/-^;*mdx* TTFs transfected with SaCas9 alone (control) or with SaCas9 plus one of the three gRNAs targeting GFP from **b**. All three gRNAs result in loss of GFP expression. GFP-gRNA-2 was selected for use in subsequent experiments due to its proximity to the colour switching mutations. GFP-gRNA-2 is referred to as GFP-gRNA in the main text. SSC, side scatter. **d**, Representative flow cytometry plots of GFP and BFP expression of GFP^+/-^;*mdx* myoblasts transfected with lipofectamine only or with SaCas9 + GFP-gRNA-2, in the absence of the BFP template. GFP^-^/BFP^-^, but not BFP^+^, cells were present in cultures transfected with SaCas9 and gRNA, consistent with NHEJ-mediated disruption without green-to-blue spectral shift.

**Fig. S2.**
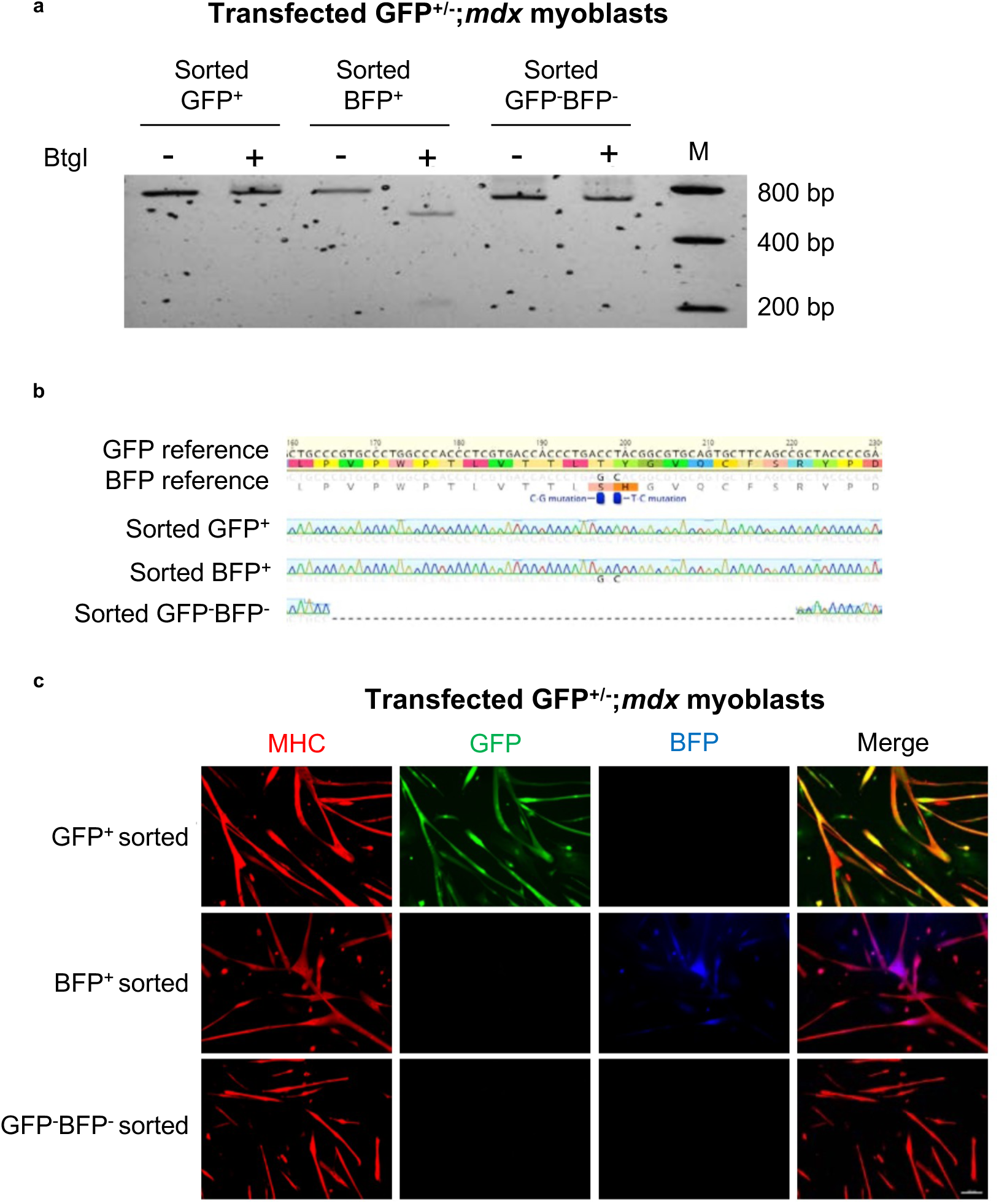
Functional and genomic confirmation of *ex vivo* edited myoblasts. **a**, Restriction fragment length polymorphism (RFLP) analysis of genomic PCR products from FACS-sorted, culture-expanded myoblasts. M, marker. **b**, Representative micrographs of myotubes differentiated from sorted GFP^+^, BFP^+^ and GFP^-^BFP^-^ myoblasts transfected with SaCas9 and GFP-gRNA-BFP-Template. Scale bar, 100 µm. Red, myosin heavy chain (MHC); Green, GFP; Blue, BFP. **c**, Sanger sequencing of PCR-amplified genomic amplicons from sorted GFP^+^, BFP^+^ and GFP^-^/BFP^-^ myoblasts transfected with SaCas9 and GFP-gRNA-BFP-Template.

**Fig. S3.**
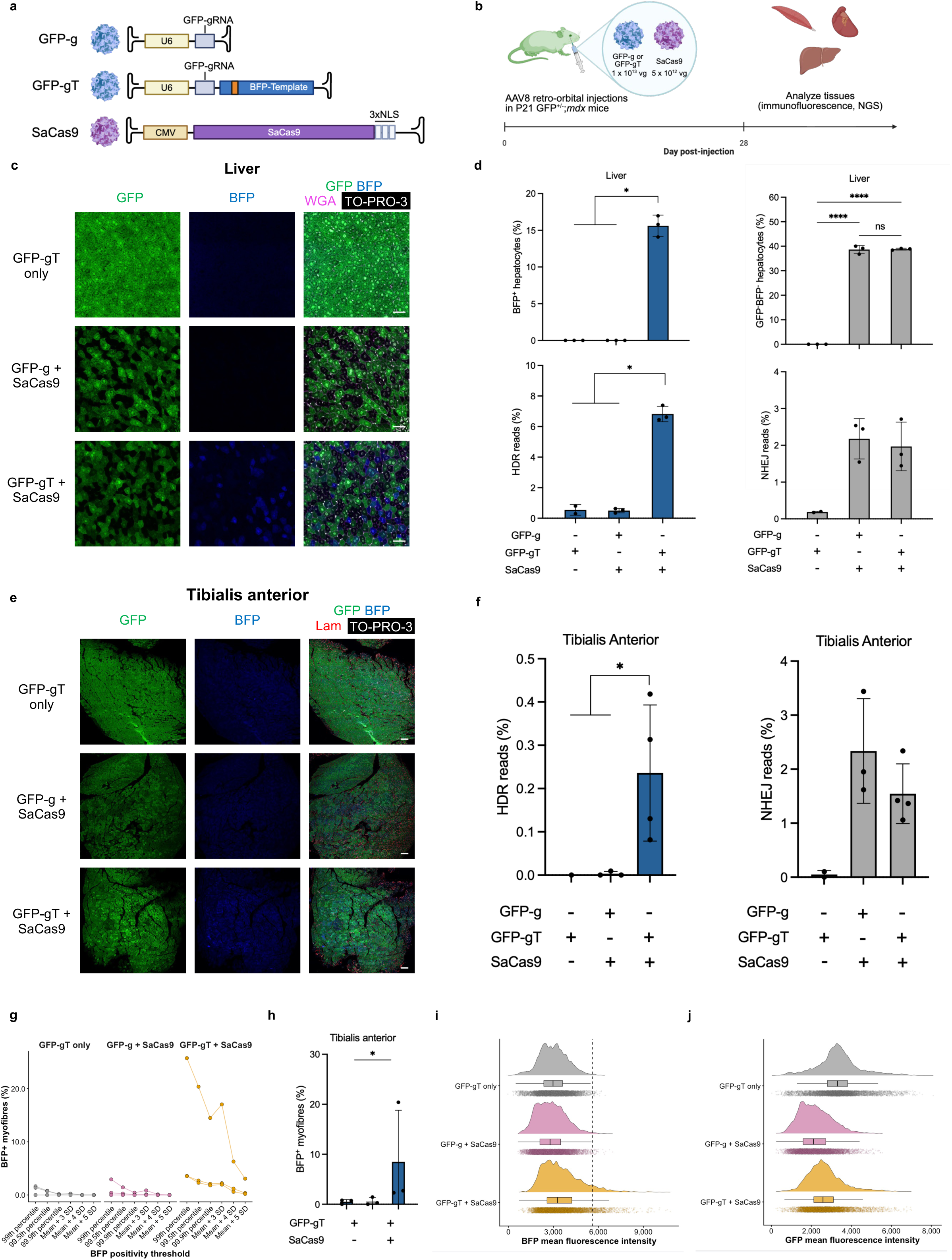
CRISPR-induced indels alone do not induce blue-shifted fluorescence in vivo. **a,**Adeno**-**associated virus constructs used for in vivo colour switching. **b**, Experimental design. Transgenic P21 GFP^+/-^; *mdx* mice carrying a single CAG-GFP allele were injected with AAV-GFP-gRNA-BFP-Template only (GFP-gT only), AAV-GFP-gRNA plus AAV-SaCas9 (GFP-g + SaCas9, or AAV-GFP-gRNA-BFP-Template plus AAV-SaCas9 (GFP-gT + SaCas9). Organs were harvested at day 28 post-injection for immunofluorescence and genomic analyses. As an additional control for batch effects, all AAV for these experiments was produced by an independent viral vector core compared to the AAV used for P21 GFP^+/-^;*mdx* experiments in Fig. 2. **c, e**, Representative immunofluorescence micrographs of liver (**c**) and tibialis anterior muscle (TA, **e**) demonstrating loss of GFP but no gain of BFP in the cutting-only group. **d, f**, Quantification of HDR and NHEJ events in liver (**d**) and TA muscle (**f**) by deep amplicon sequencing. For HDR and BFP^+^ quantification, samples from GFP-gT and GFP-gT + SaCas9 were pooled prior to statistical testing. Samples with aligned reads < 1,000 were excluded from quantification. Conditions with fewer than two biological replicates after quality control are shown descriptively and were not included in statistical testing. **g**, Quantification of BFP-positive myofibres at different mean fluorescence intensity thresholds. Each set of connected points represents one biological replicate. The frequency of GFP- myofibres were not quantified in skeletal muscle myofibres due to the high degree of multinucleation in this tissue, which prevents detection of complete loss of GFP fluorescence unless all myonuclei contributing to a myofibre are edited. To avoid pseudoreplication, myofibres from only a single cryosection per biological replicate were included. Samples from mice injected with GFP-gT alone or GFP-g and SaCas9 show minimal BFP positivity, while those injected with GFP-gT and SaCas9 show a substantial BFP-positive fraction. **h**, Quantification of BFP^+^ myofibres. **i, j**, Raincloud plots comparing the mean fluorescence intensity of BFP and GFP in GFP-gT only, GFP-g + SaCas9 and GFP-gT + SaCas9 injected mice. Data are presented as mean ± SD. **d** and **f** were analysed with the Mann-Whitney *<u>U</u>* test for HDR and BFP quantification or one-way ANOVA with Tukey’s multiple comparisons for NHEJ. **h** was analysed with one-way ANOVA with Tukey’s multiple comparisons. \**P* < .05, \*\*\*\**P* < .0001. n.s., not significant. GFP-gT, n = 3; GFP-g + SaCas9, n = 3; GFP-gT + SaCas9, n = 4.

**Fig. S4.**
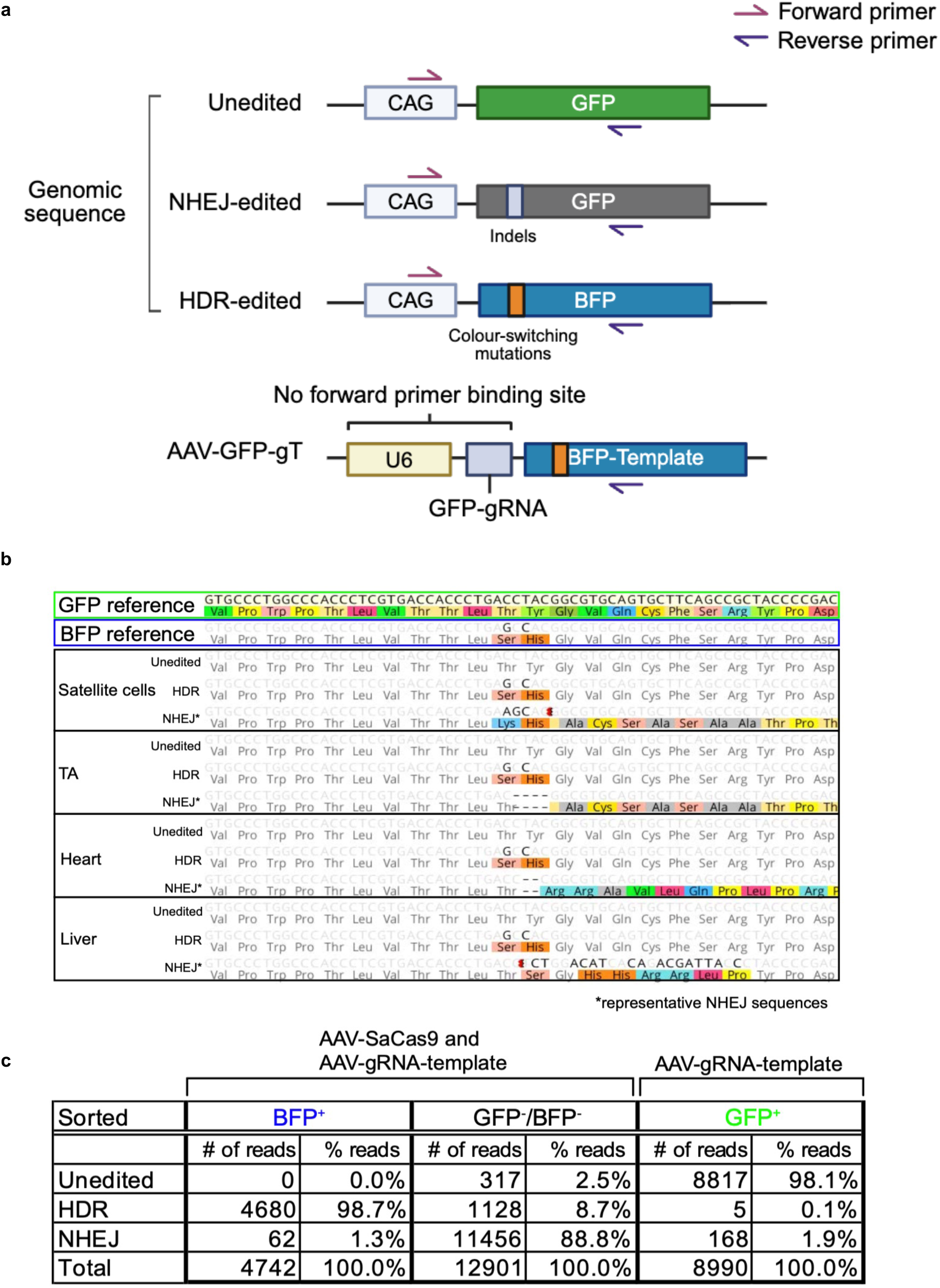
Deep amplicon sequencing validates in vivo NHEJ and HDR editing events. **a**, Schematic of genomic locus, BFP donor template, and primer design used for amplification of edited alleles. The forward primer binds upstream of the GFP/BFP start site on the genomic sequence, but not template DNA, while the reverse primer binds downstream of the Cas9 cutting site and colour-switching substitutions. **b**, Representative aligned sequences from deep amplicon sequencing of unedited, HDR-edited, and NHEJ-edited alleles from satellite cells, tibialis anterior (TA) muscle, heart, and liver tissues of P21 AAV-GFP-to-BFP injected GFP^+/-^;*mdx* mice. *representative NHEJ variants are shown. **c**, Summary table of deep sequencing read counts and allele frequences corresponding to unedited, HDR, and NHEJ editing detected in FACS-sorted satellite cells from P21 GFP^+/-^;*mdx* mice injected with AAV-GFP-gRNA-Template (GFP-gT only) or AAV-SaCas9 and AAV-gRNA-BFP-Template (GFP-gT + SaCas9). BFP^+^ and GFP^-^BFP^-^ cells were sorted from AAV-SaCas9 and AAV-gRNA-BFP-Template injected experimental mice, while GFP^+^ cells were sorted from AAV-gRNA-BFP-Template injected control mice. GFP-gT, n = 3; GFP-gT + SaCas9, n = 4.

**Fig. S5.**
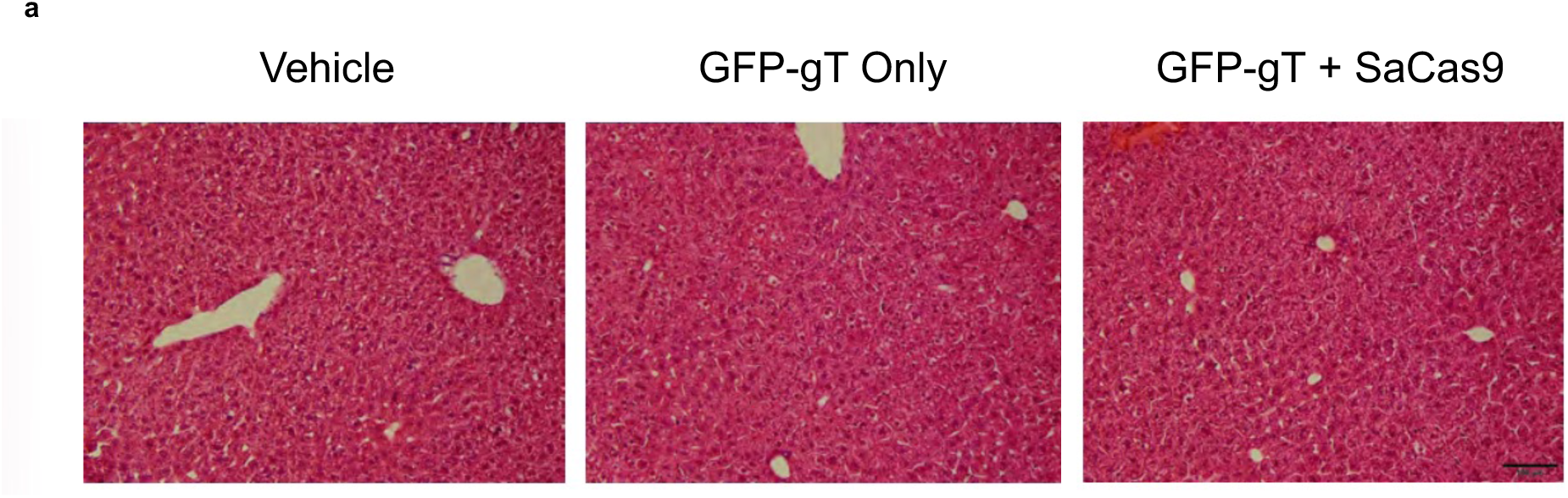
Systemic high-dose AAV delivery does not result in histological evidence of hepatotoxicity. **a,**Representative hematoxylin and eosin (H&E) staining of liver cryosections from GFP^+/-^;*mdx* mice injected systemically with vehicle, AAV-GFP-gRNA-BFP-Template (GFP-gT), or AAV-GFP-gRNA-Template (GFP-gT) and AAV-SaCas9 (SaCas9). Scale bar, 100 µm.\

**Fig. S6.**
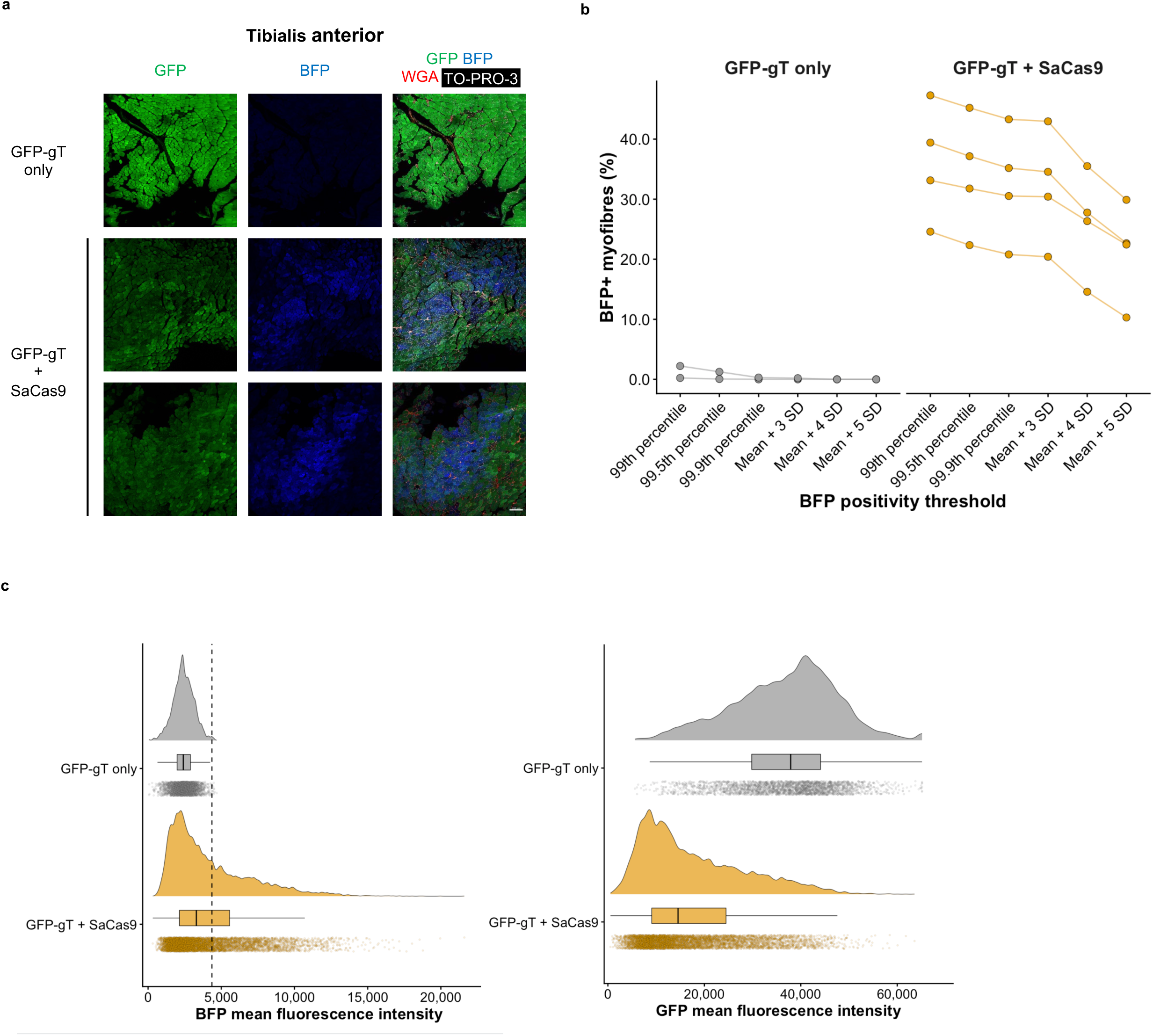
Whole-section imaging of tibialis anterior muscle from GFP^+/-^;*mdx* mice injected with AAV-GFP-to-BFP and threshold-based identification of BFP-positive myofibres. **a,**Representative stitched immunofluorescence micrographs of whole tibialis anterior muscle cryosections from mice receiving AAV-GFP-gRNA-Template (GFP-gT) or GFP-gT together with AAV-SaCas9 (SaCas9). Green, GFP; Blue, BFP; Red, Wheat germ agglutinin (WGA); white, TO-PRO-3. Scale bar, 200 µm. **b**, Quantification of BFP-positive myofibres at different mean fluorescence intensity thresholds. Each set of connected points represents one biological replicate. To avoid pseudoreplication, myofibres from only a single cryosection per biological replicate were included. Samples from mice injected with GFP-gT show minimal BFP positivity, while those injected with GFP-gT and SaCas9 show a substantial BFP-positive fraction. For quantification in Figure 2**g**, a threshold based on the 99.5^th^ percentile of fibre BFP mean fluorescence intensity in negative controls was used. **c**, Raincloud plots comparing the mean fluorescence intensity of BFP and GFP in GFP-gT only and GFP-gT + SaCas9 injected mice. GFP-gT only, n = 3; GFP-gT + SaCas9, n = 4.

**Fig. S7.**
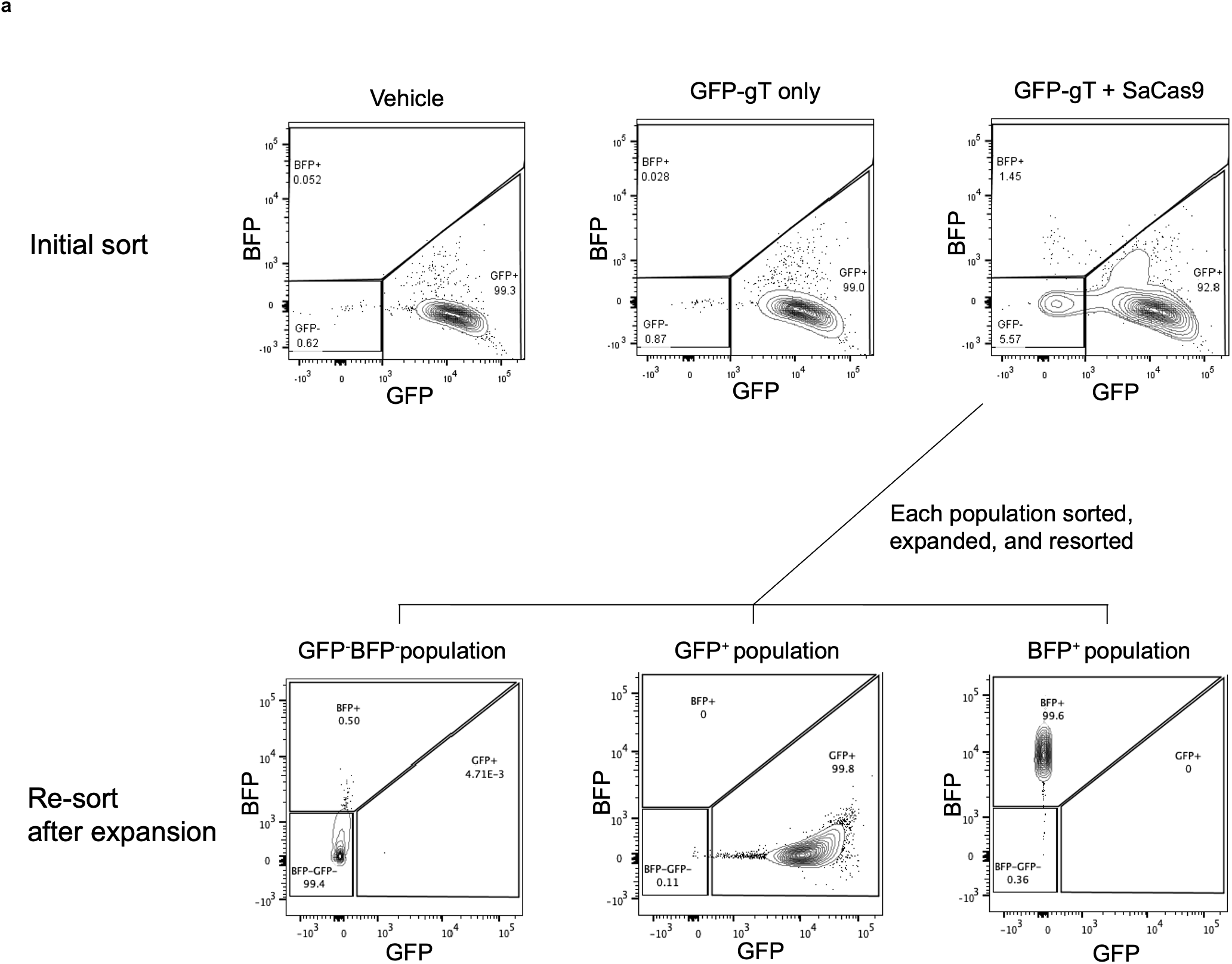
Fluorescence stability of in vivo CRISPR-NHEJ and CRISPR-HDR edited muscle stem cells. **a**, Representative flow cytometry plots showing analysis of GFP and BFP expression by skeletal muscle satellite cells isolated from juvenile GFP^+/-^;*mdx* mice previously injected intravenously with vehicle AAV-GFP-gRNA-BFP-Template alone (GFP-gT only) as control or AAV-GFP-gRNA-BFP-Template and AAV-SaCas9 (GFP-gT + SaCas9). FACS-sorted populations were expanded separately in culture for 2 weeks and then harvested for re-analysis (shown in **b**). **b**, Representative flow cytometric analysis of GFP and BFP expression in culture-expanded GFP^-^BFP^-^, GFP^+^, and BFP^+^ cells previously sorted from AAV-HDR injected mice. GFP-gT, n = 3; GFP-gT + SaCas9, n = 4.

**Figure S8.**
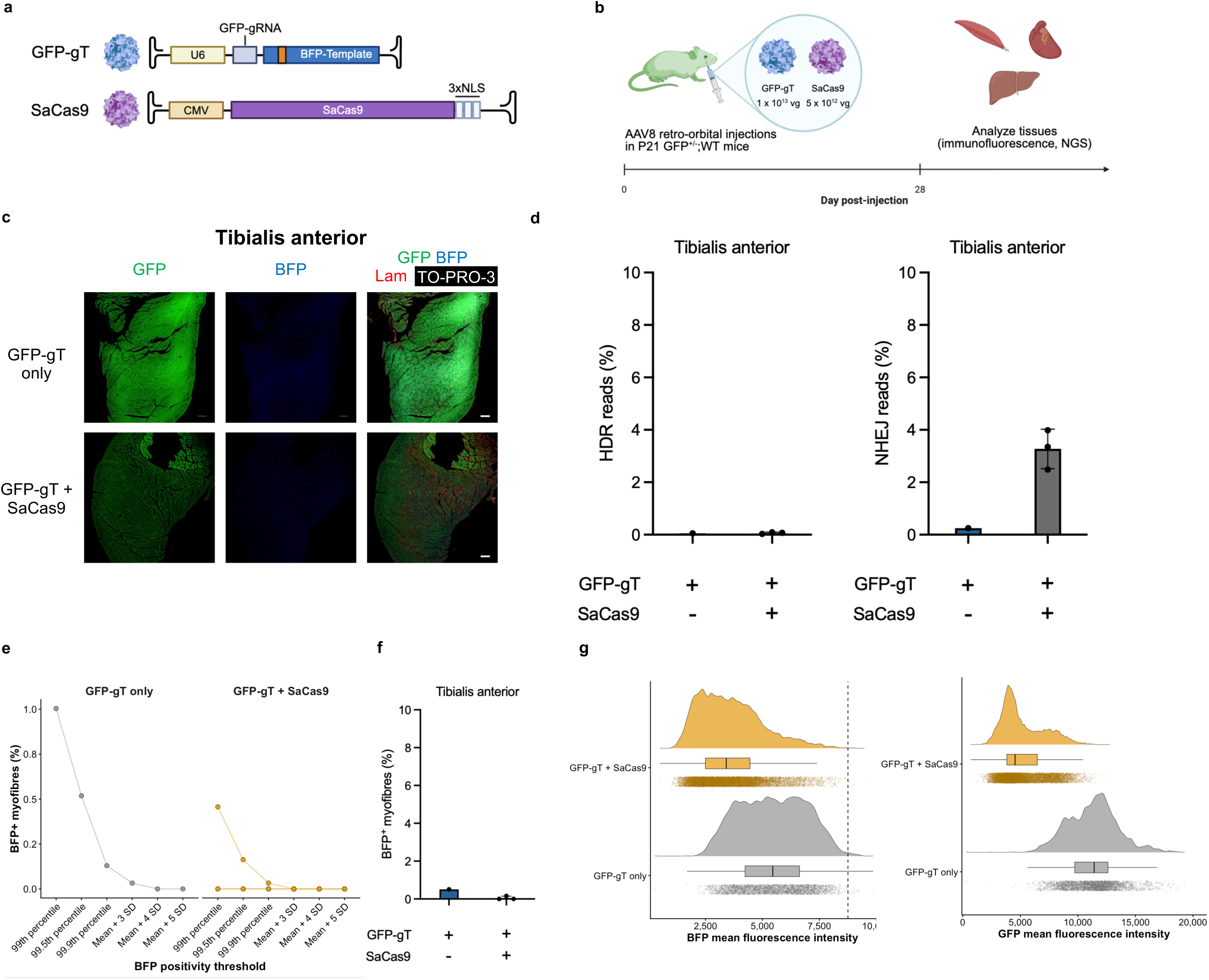
Wild-type non-dystrophic skeletal muscle restricts detectable in vivo CRISPR-HDR . **a**, Adeno-associated virus constructs used for in vivo colour switching. **b**, Experimental design. Transgenic P21 GFP**^+/-^**;WT mice carrying a single CAG-GFP allele were injected with AAV-GFP-gRNA-BFP-Template only (GFP-gT) or AAV-GFP-gRNA-BFP-Template (GFP-gT) plus AAV-SaCas9 (SaCas9). **c**, Immunofluorescence micrographs of tibialis anterior muscle (TA) Green, GFP; Blue, BFP; Red, laminin; White, TO-PRO-3. **d**, Quantification of HDR and NHEJ events in TA muscle by deep amplicon sequencing. **e**, Quantification of BFP-positive myofibres at different mean fluorescence intensity thresholds. Each set of connected points represents one biological replicate. To avoid pseudoreplication, myofibres from only a single cryosection per biological replicate were included. Samples from mice injected with GFP-gT alone or GFP-gT and SaCas9 show minimal BFP positivity**. f**, Quantification of BFP^+^ cells in TA muscle. **g**, Raincloud plots comparing the mean fluorescence intensity of BFP and GFP in GFP-gT only and GFP-gT + SaCas9 injected mice. GFP-gT only, n = 1; GFP-gT + SaCas9, n = 3. Due to limited sample size, data are presented descriptively and no statistical tests were performed.

**Figure S9.**
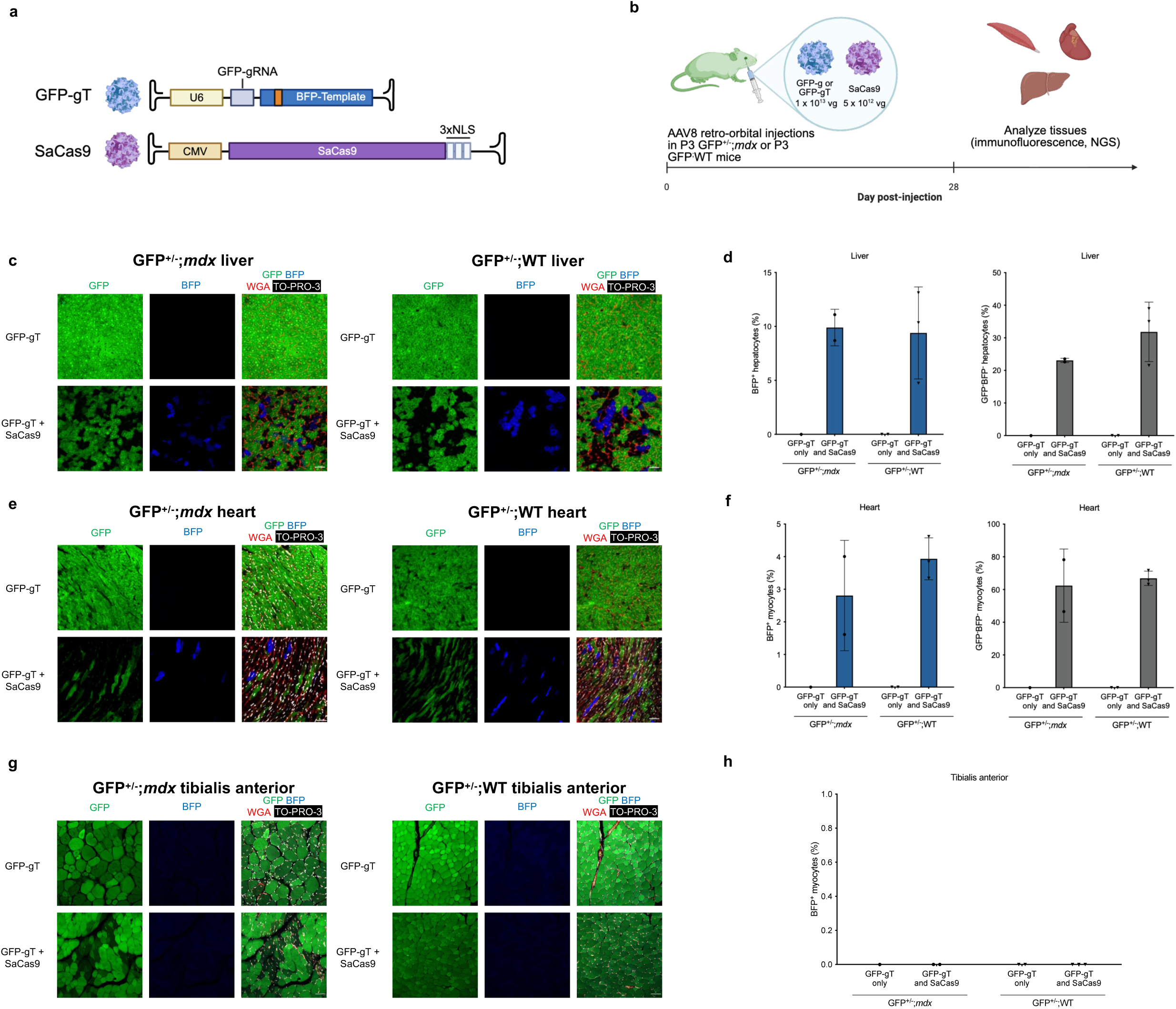
Early neonatal (P3) delivery of systemic AAV-GFP-to-BFP reveals distinct developmental capacities for HDR. **a**, Adeno-associated virus constructs used for in vivo colour switching. **b**, Experimental design. Transgenic P3 GFP^+/-^;*mdx* or GFP^+/-^;WT mice carrying a single CAG-GFP allele were injected with AAV-GFP-gRNA-BFP-Template only (GFP-gT) or AAV-GFP-gRNA-BFP-Template (GFP-gT) plus AAV-SaCas9 (SaCas9). **c**, **d,** Representative immunofluorescence micrographs (**c**) and quantification (**d**) of live cryosections; Green, GFP; Blue, BFP; Red, laminin; White, TO-PRO-3. and quantification of liver cryosections. **e**,**f,** Representative immunofluorescence micrographs (e) and quantification (f) of tibialis anterior (TA); Green, GFP; Blue, BFP; Red, laminin; White, TO-PRO-3. **g,h**, Representative immunofluorescence micrographs (**g**) of the tibialis anterior (TA) muscle. Data are shown as mean ± SD. n = 5 for experimental groups (n = 2 GFP^+/-^;*mdx*, n = 3 GFP^+/-^;WT), n = 3 for control groups (n = 1 GFP^+/-^;*mdx*, n = 2 GFP^+/-^;WT). Due to limited sample size, data are presented descriptively and no statistical tests were performed.

**Figure S10.**
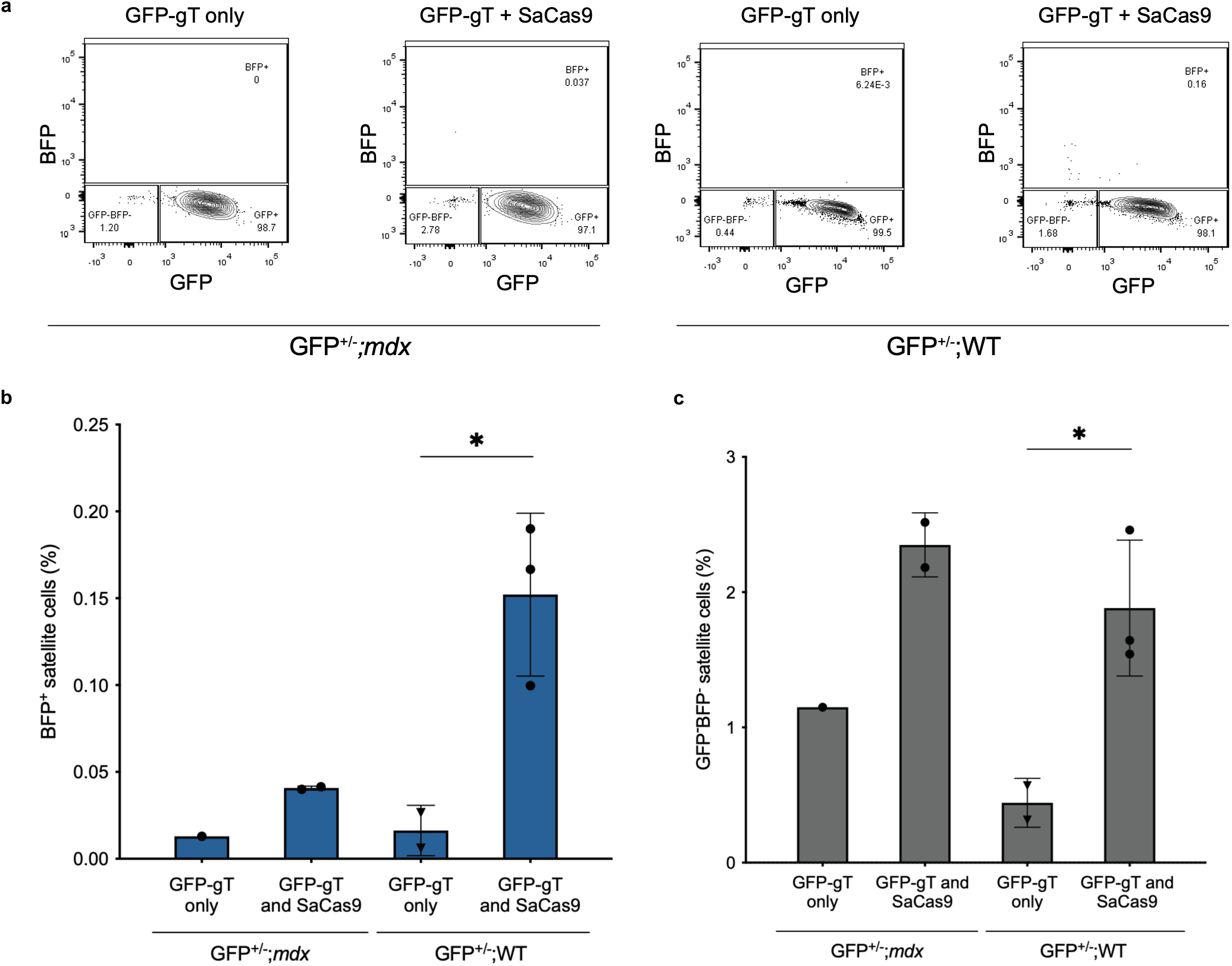
Satellite cells in neonatal (P3) skeletal muscles are infrequently targeted with systemic AAV-GFP-to-BFP. **a,**Representative flow cytometric analysis of skeletal muscle satellite cells isolated from neonatal (P3) GFP^+/-^;*mdx* and GFP^+/-^;WT mice 28 days after intraperitoneal injection with AAV-GFP-gRNA-BFP-Template (GFP-gT) alone as control or with AAV-GFP-gRNA-BFP-Template and AAV-SaCas9 (GFP-gT + SaCas9). **b, c** Quantification of in vivo edited BFP^+^ satellite cells (**b**) and GFP^-^BFP^-^ satellite cells (**c**). Data are shown as mean ± SD. \**P* < .05, one-way analysis of variance (ANOVA) with Tukey’s multiple comparisons test. n = 5 for experimental groups (n = 2 GFP^+/-^;*mdx*, n = 3 GFP^+/-^;WT), n = 3 for control groups (n = 1 GFP^+/-^;*mdx*, n = 2 GFP^+/-^;WT).

**Figure S11.**
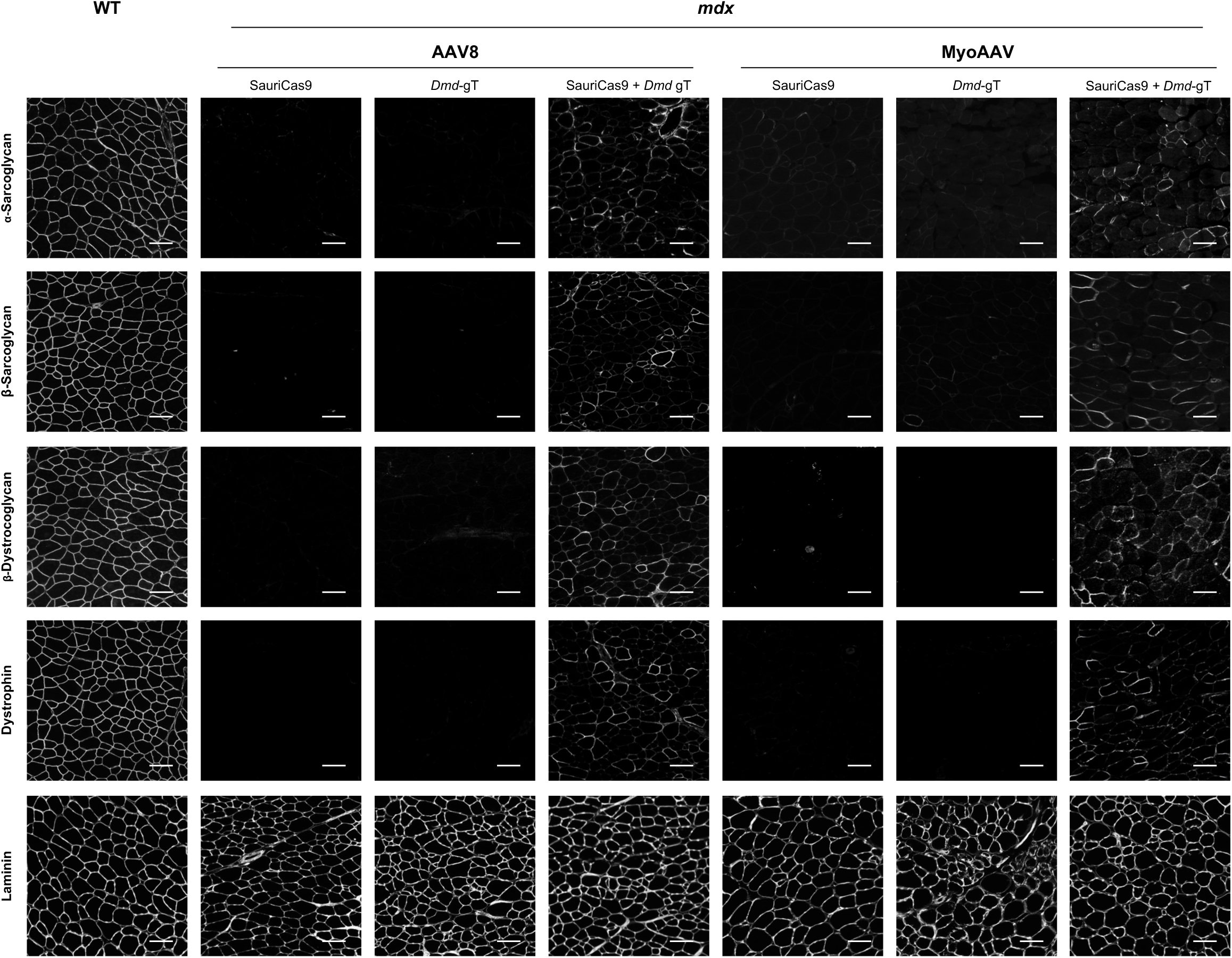
Systemic dual delivery of AAV-SauriCas9 and AAV-*mdx*-gRNA-*Dmd*-Template restores the dystrophin-associated protein complex (DAPC). Immunofluorescence staining for the indicated dystrophin-associated protein complex (DAPC) components in muscle sections from wild-type mice (WT, left) or *mdx* mice injected retro-orbitally with AAV-SauriCas9 (SauriCas9), AAV-*mdx*-gRNA-*Dmd*-Template (*Dmd*-gT), AAV-SauriCas9 plus AAV-*mdx*-gRNA-*Dmd*-Template (SauriCas9 + *Dmd*-gT), MyoAAV-SauriCas9 (SauriCas9), MyoAAV-*mdx*-gRNA-*Dmd*-Template (*Dmd*-gT), or MyoAAV-SauriCas9 plus MyoAAV-*mdx*-gRNA-*Dmd*-Template (SauriCas9 + *Dmd*-gT). Scale bar = 100 µm.

**Figure S12.**
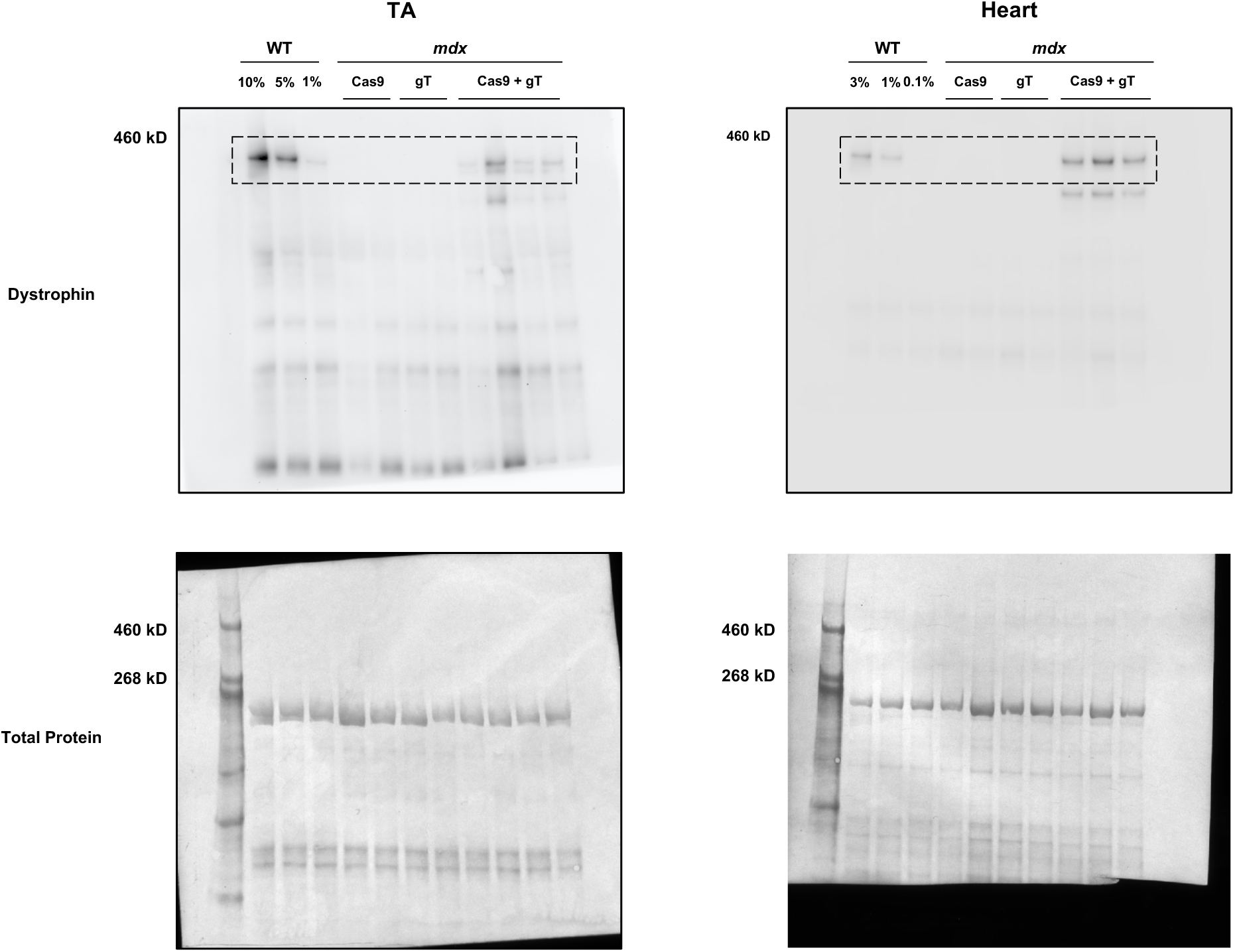
Systemic dual delivery of AAV-SauriCas9 and AAV-*mdx*-gRNA-*Dmd*-Template restores dystrophin protein expression in *mdx* mice. Full membrane images of dystrophin protein expression detected by Western Blot of protein lysate from TA (left) or heart (right) of the indicated wild-type (WT) mice or *mdx* mice injected with AAV-SauriCas9 (SauriCas9), AAV-*mdx*-gRNA-*Dmd*-Template (*Dmd*-gT), or AAV-SauriCas9 plus AAV-*mdx*-gRNA-*Dmd*-Template (SauriCas9 + *Dmd*-gT). To estimate the efficiency of dystrophin protein rescue, the first 3 lanes were loaded with different percentages (10% - 5% - or 1% for TA and 3% - 1% - or 0.1% for heart) of WT lysate. Coomassie blue staining of each membrane is shown below each blot, demonstrating equivalent protein loading. Dashed squares highlight the regions of each blot used for quantification of dystrophin protein levels (427 kDa).

**Figure S13.**
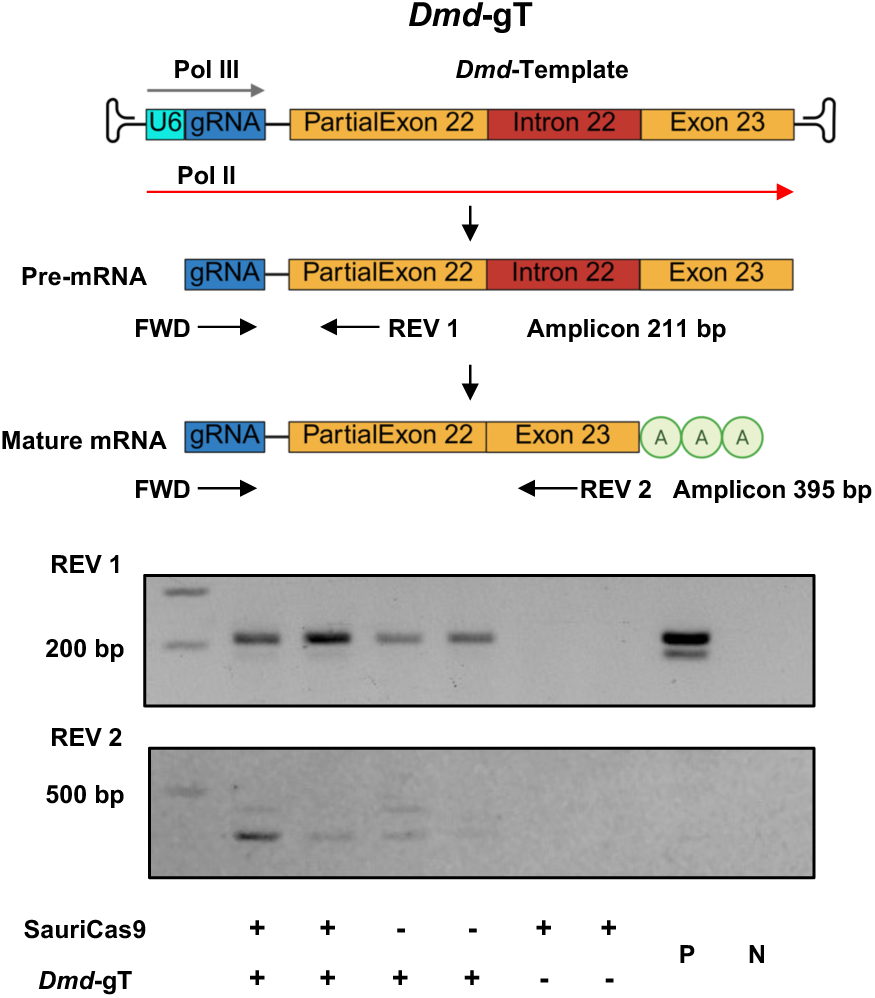
AAV-*mdx*-gRNA-*Dmd*-Template injected mice transcribes a transcript that contains *mdx*-gRNA and *Dmd*-template sequences. The schematic demonstrates how the RNA polymerase II (Pol II) activity of U6 promoter could transcribe a transcript that contains *mdx*-gRNA and *Dmd*-template sequences. To test this hypothesis, oligo-dT reverse transcribed heart cDNA of *mdx* mice injected with the indicated AAVs, *mdx*-gRNA-*Dmd*-Template AAV transfer plasmid (P), or water (N) were used as templates. Forward primer targeting the *mdx*-gRNA locus (FWD) and reverse primer 1 targeting *Dmd* exon 22 (REV 1) were used to confirm the presence of *mdx*-gRNA and *Dmd*-Template sequences containing transcripts. Amplicon size is expected to be 211 base pair (bp). FWD and reverse primer 2 targeting *Dmd* exon 23 (REV 2) were used to confirm the excision of *Dmd* intron 22 in the transcript that contains *mdx*-gRNA and *Dmd*-Template. If *Dmd* intron 22 is excised, amplicon size is expected to be 395 bp.

**Figure S14.**
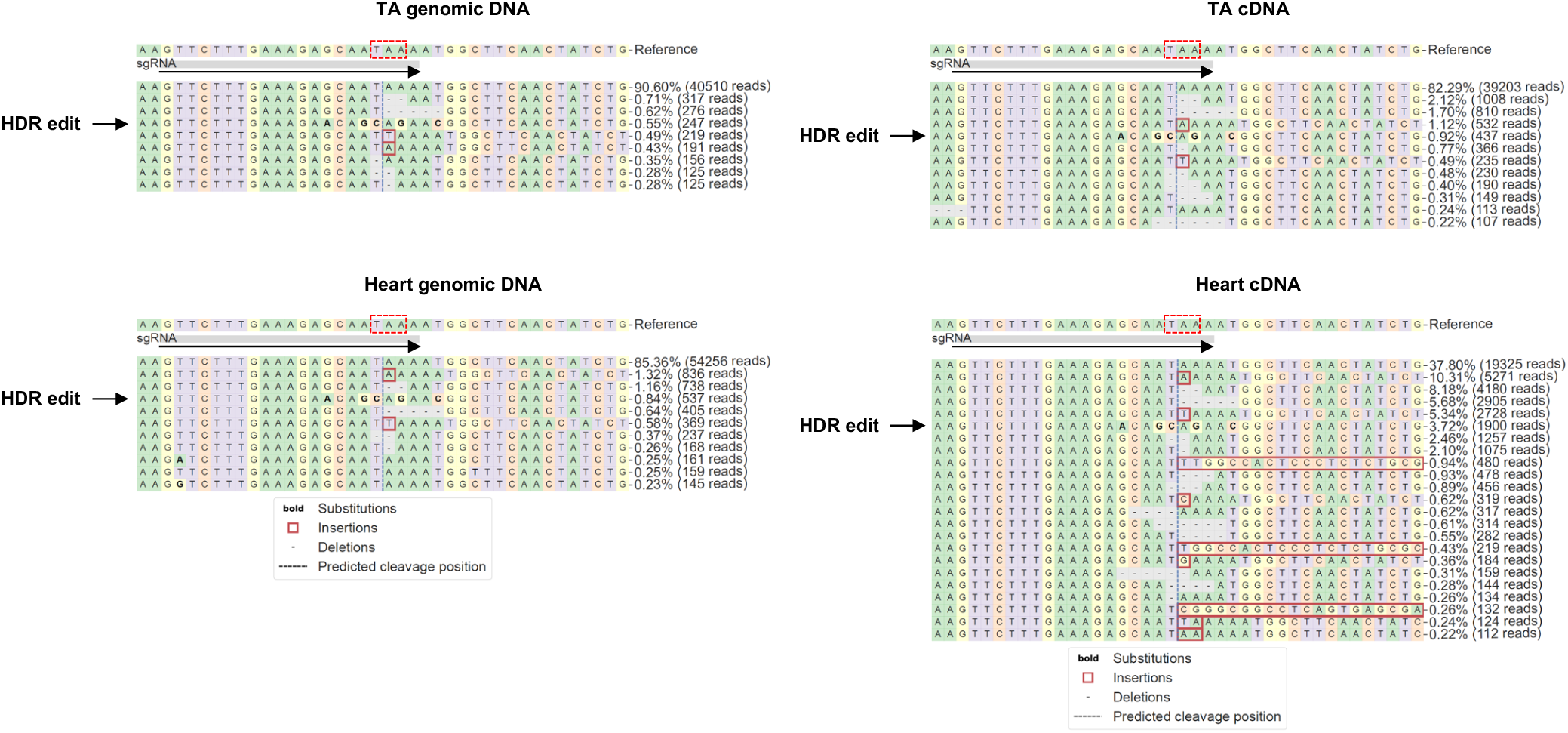
Amplicon sequencing analysis for AAV-SauriCas9 and AAV-*mdx*-gRNA-*Dmd*-Template mediated editing outcomes at the *mdx* nonsense mutation and gRNA binding site. TA and heart cDNA and genomic were used for amplicon sequencing. Sequencing results were analyzed with CRISPREsso2 algorithm. Representative allele frequency tables are shown. Dashed red box indicates the nonsense mutation stop codon, and arrow indicates the orientation of gRNA binding. Reads bearing HDR edits are highlighted with black arrows.

**Figure S15.**
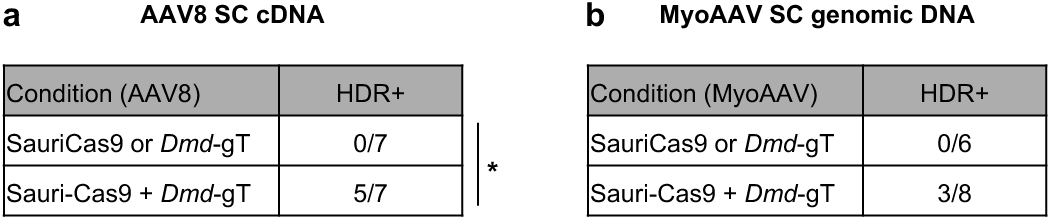
Systemic dual delivery of AAV-SauriCas9 and AAV-*mdx*-gRNA-*Dmd*-Template corrects *mdx* mutation in satellite cells. Satellite cells from control and experimental mice described in Fig. 3 and Fig. 4 were FACS-sorted. **a**, Satellite cells isolated from mice used in Fig. 3 were expanded *in vitro* for 4 days. cDNA from these satellite cells were used for amplicon sequencing at the *mdx* mutation site. **b**, Genomic DNA from satellite cells isolated from mice used in Fig. 4 were used for amplicon sequencing at the *mdx* mutation site. The tables indicate the number of mice with the indicated condition in which HDR edits were detected (low rates, < 0.6%). * indicates *P* < .05, analyzed with two-tailed Fisher’s exact test.

**Figure S16.**
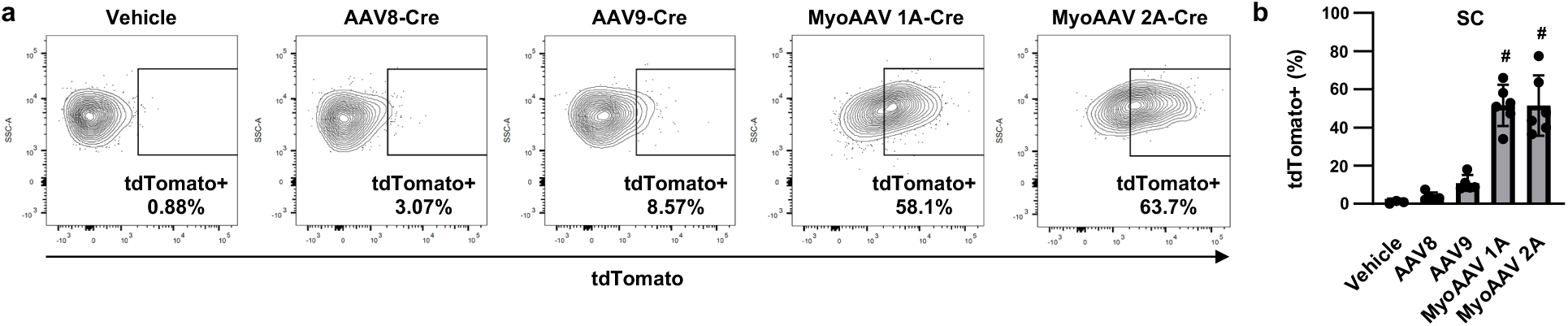
Superior transduction of muscle satellite cells by MyoAAV vectors. **a**, Representative flow cytometry plots of satellite cells isolated from juvenile loxp-STOP-loxp-tdTomato transgenic mice (*Ai14*) mice retro-orbitally injected with vehicle, or 1 x 10^12^ vg/kg of AAV8, AAV9, MyoAAV1A, or MyoAAV2A carrying Cre endonuclease 4 days after injection to analyze for tdTomato expression, induced by Cre activity and indicative of AAV transduction. **b**, Summary of % tdTomato+ satellite cells isolated from *Ai14* mice treated with each indicated AAV serotype driving Cre endonuclease, representing AAV transduction, vehicle, n = 3, AA8 and AAV9, n = 4, and MyoAAV 1A and 2A, n = 5, data is presented as mean ± SD. **b** is analyzed with one-way ANOVA, # indicates *P* < .001 compared to vehicle, AAV8, and AAV9 conditions.

**Figure S17.**
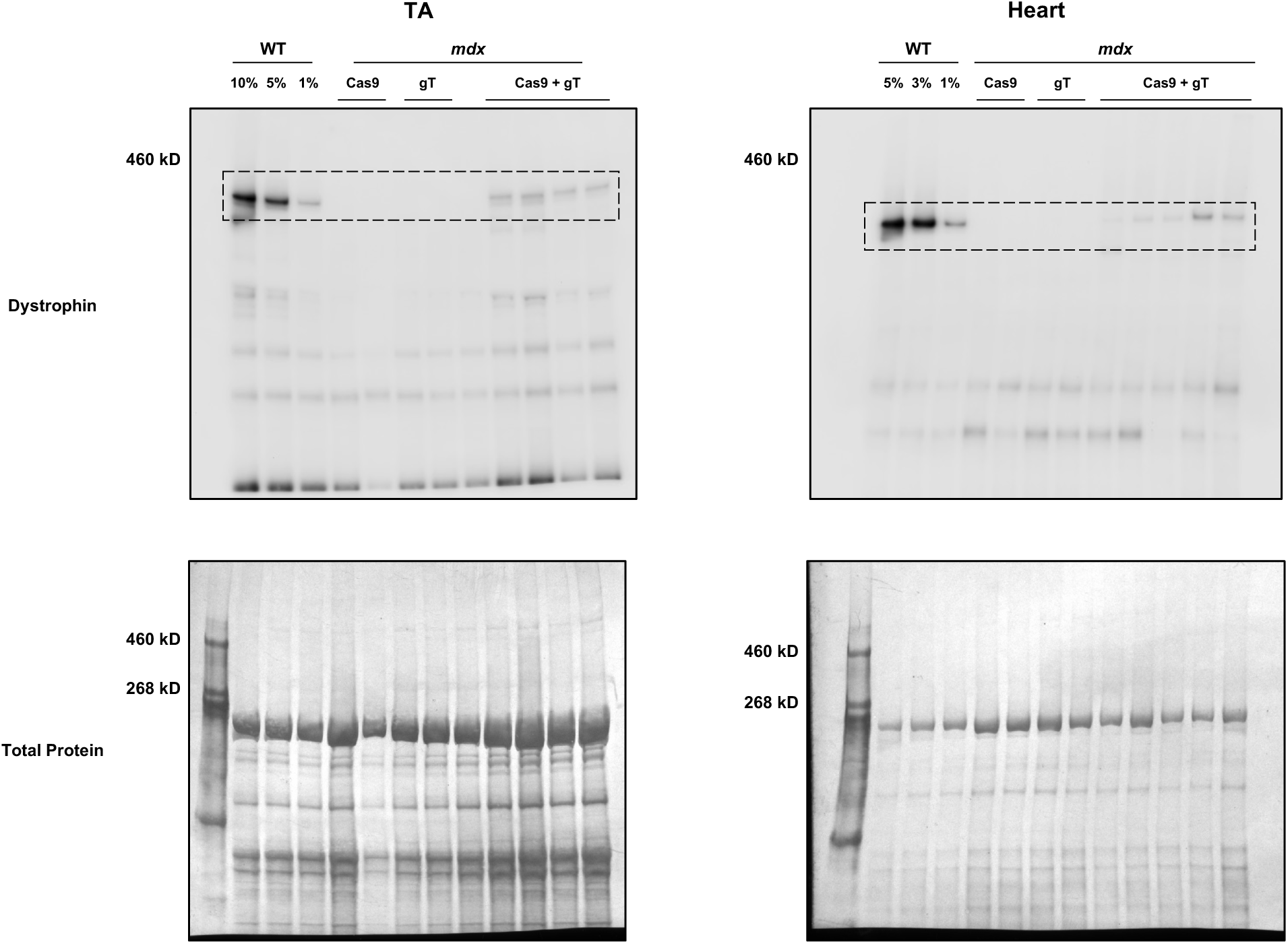
Systemic dual delivery of MyoAAV-SauriCas9 and MyoAAV-*mdx*-gRNA-*Dmd*-Template restores dystrophin protein expression in *mdx* mice. Full membrane images of dystrophin protein expression detected by Western Blot of protein lysate from TA (left) or heart (right) of the indicated wild-type (WT) mice, or *mdx* mice injected with MyoAAV-SauriCas9 (SauriCas9), MyoAAV-*mdx*-gRNA-*Dmd*-Template (*Dmd*-gT), or AAV-SauriCas9 plus MyoAAV-*mdx*-gRNA-*Dmd*-Template (SauriCas9 + *Dmd*-gT). To estimate the efficiency of dystrophin protein rescue, the first 3 lanes were loaded with different percentages (10% - 5% - or 1% for TA and 5% - 3% - or 1% for heart) of WT lysate. Coomassie blue staining of each membrane is shown below each blot, demonstrating equivalent protein loading. Dashed squares highlight the regions of each blot used for quantification of dystrophin protein levels (427 kDa).

**Figure S18.**
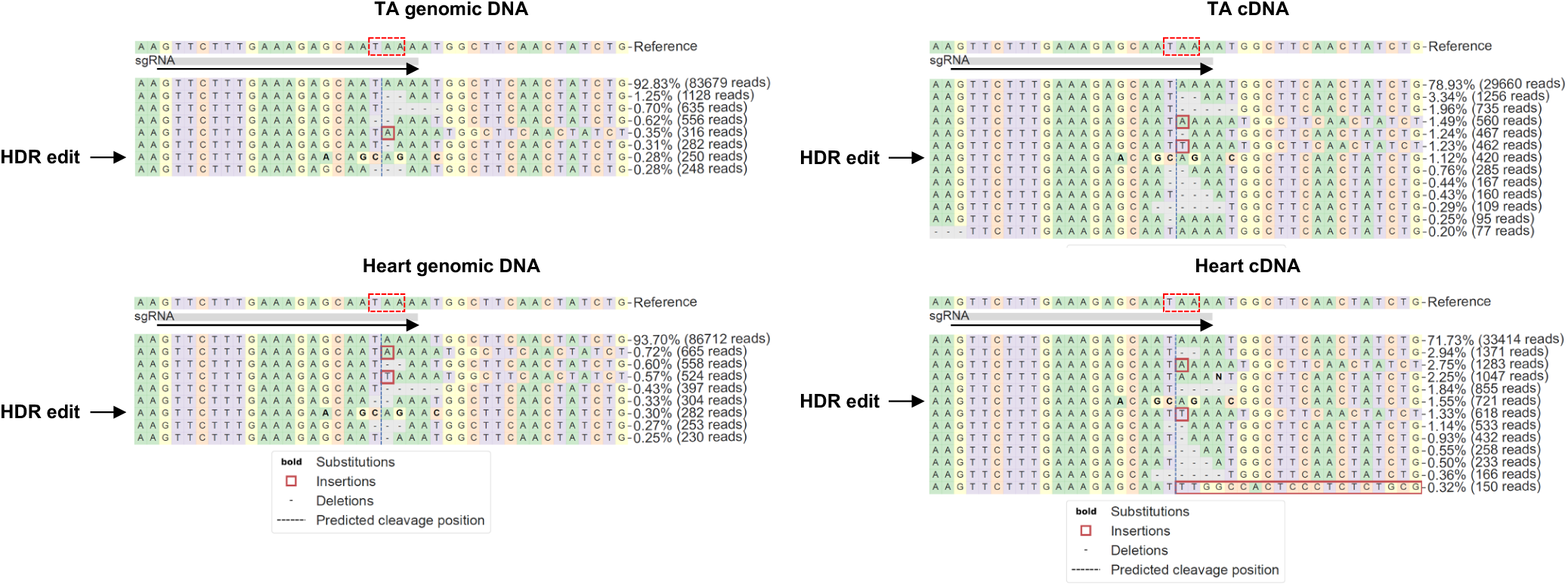
Amplicon sequencing analysis for MyoAAV-SauriCas9 and MyoAAV-*mdx*-gRNA-*Dmd*-Template mediated editing outcomes at the *mdx* nonsense mutation and gRNA binding site. TA and heart cDNA and genomic were used for amplicon sequencing. Sequencing results were analyzed with CRISPREsso2 algorithm. Representative allele frequency tables are shown. Dashed red box indicates the nonsense mutation stop codon, and arrow indicates the orientation of gRNA binding. Reads bearing HDR edits are highlighted with black arrows.

